# AI-Driven Variant Annotation for Precision Oncology in Breast Cancer

**DOI:** 10.1101/2025.03.14.643357

**Authors:** Kriti Shukla, Yue Wang, Philip M. Spanheimer, Elizabeth Brunk

**Author notes:** Correspondence should be addressed to: Elizabeth Brunk.

## Abstract

Interpreting the functional impact of genomic variants remains a major challenge in precision oncology, particularly in breast cancer, where many variants of unknown significance (VUS) lack clear therapeutic guidance. Current annotation strategies focus on frequent driver mutations, leaving rare or understudied variants unclassified and clinically uninformative. Here, we present an AI/ML-driven framework that systematically identifies variants associated with key breast cancer phenotypes, including ESR1 and EZH2 activity, by integrating genomic, transcriptomic, structural, and drug response data. Using DepMap and TCGA datasets, we analyzed >12,000 variants across breast cancer genomes, identifying structurally clustered mutations that share functional consequences with well-characterized oncogenic drivers. This approach reveals that mutations in PIK3CA, TP53, and other genes strongly associate with ESR1 signaling, challenging conventional assumptions about endocrine therapy response. Additionally, EZH2-associated variants emerge in unexpected genomic contexts, suggesting new targets for epigenetic therapies. By shifting from frequency-based to structure-informed classification, we expand the set of potentially actionable mutations, enabling improved patient stratification and drug repurposing strategies. This work provides a scalable, clinically relevant method to accelerate variant annotation, offering new insights into drug sensitivity and resistance mechanisms. Future validation efforts will refine these predictions and integrate clinical outcomes to guide personalized treatment strategies. Our findings highlight the transformative potential of AI/ML in redefining cancer variant interpretation, bridging the gap between genomics, functional biology, and precision medicine.

**Study Highlights:** *Current Knowledge:* Breast cancer treatment decisions are increasingly guided by genomic profiling, yet most clinical actionability is based on frequent driver mutations (e.g., PIK3CA, TP53, ESR1). Many variants of unknown significance (VUS) remain unclassified, and current annotation methods are slow, relying on manual curation or low-throughput assays, leaving rare mutations uncharacterized.

*Study Focus:* This study applies AI/ML-driven variant annotation to systematically identify mutations that drive key breast cancer phenotypes, such as ESR1 and EZH2 activity, beyond currently known mutations. By using structural and functional clustering, we assess whether rare and understudied mutations can be prioritized for clinical relevance.

**Key Findings:**

● Analyzed >12,000 variants across breast cancer genomes, integrating multi-omic and structural data.
● Identified strong ESR1-associated mutations in PIK3CA, TP53, and other genes, expanding the landscape of actionable mutations.
● Discovered EZH2-associated variants in unexpected contexts, revealing potential epigenetic therapy targets.
● Demonstrated that spatial clustering of mutations within proteins predicts functional consequences, even for rare mutations.

**Clinical and Translational Impact:**

● Scalable AI-powered framework accelerates variant annotation and functional classification.
● Enables faster identification of actionable mutations and improves patient stratification for targeted therapies.
● Provides a data-driven approach to refine clinical trial design, expanding therapy options for patients lacking clear genomic-based treatment guidance.

## Introduction

Genomic profiling plays an essential role in breast cancer treatment selection, particularly in determining targeted therapies for hormone receptor-positive tumors. Patients with estrogen receptor-positive (ER+) breast cancer are treated with endocrine therapy(1,2), including tamoxifen, aromatase inhibitors (e.g., letrozole, anastrozole, exemestane) and selective estrogen receptor degraders (e.g., fulvestrant), which suppress estrogen-estrogen receptor signaling (3–5). However, responses remain variable(6), with *de novo* and acquired resistance leading to many treatment failures(7). While some ESR1 mutations (e.g., Y537S, D538G) are known to drive endocrine resistance(8), this is not the cause in most patients and the underlying genomic factors contributing to resistance for many patients remain unclear(6). Beyond endocrine therapy, targeted approaches such as PI3K, AKT, and mTOR inhibitors have demonstrated efficacy in patients with specific genomic alterations(9). Yet, not all patients with these mutations respond uniformly, and many patients lack clearly actionable variants(9).

A major barrier to optimizing treatment stratification is the high number of variants of unknown significance (VUS) in breast cancer(10), particularly those that may influence ESR1 signaling, epigenetic regulation, and drug response. While frequent mutations in BRCA1/2, TP53, and PIK3CA are well-characterized(11–14), many low-frequency or previously unannotated variants remain poorly understood, leaving clinicians without clear guidance on how they influence treatment decisions(15). Current annotation strategies rely on slow, manual curation methods or low-throughput functional assays, which focus predominantly on frequent driver mutations. As a result, patients who lack well-characterized mutations are often treated with conventional therapies(16), potentially missing opportunities for more precise, biomarker-driven treatment selection.

To address this, we need scalable, high-throughput approaches to systematically classify variants based on functional impact rather than occurrence frequency. The ability to identify and prioritize mutations that drive key breast cancer phenotypes, such as estrogen signaling or epigenetic reprogramming, could significantly refine treatment strategies. By leveraging AI/ML (Artificial Intelligence/ Machine Learning) and multi-omic data resources such as DepMap (Dependency Map)(17,18) and TCGA (The Cancer Genome Atlas)(19), we can systematically link genomic variants to molecular phenotypes, including transcriptional activity, protein dependencies, and drug response profiles. This allows us to rank and prioritize variants based on not just their prevalence, but also their functional impact, druggability and clinical outcome.

Here, we present an AI/ML-based approach that identifies variant clusters strongly associated with ESR1 and EZH2 activity, two key regulators of breast cancer progression and therapy response. Our approach systematically maps mutations to 3D protein structures from AlphaFold(20), allowing us to identify clusters of mutations in close spatial proximity that are expected to share functional impact, even when some variants are rare or previously unannotated. By shifting the annotation focus from mutation frequency to structural and functional impact, we significantly expand the pool of potentially actionable variants in breast cancer.

Our findings reveal that specific mutations in PIK3CA, TP53, and other genes strongly associate with ESR1 activity, suggesting they may modulate estrogenic signaling and endocrine therapy response in unexpected ways. Similarly, we identify EZH2-associated variant clusters across multiple genes, which may have important implications for emerging epigenetic therapeutic strategies in breast cancer. Beyond these well-known drivers, our AI/ML framework also identifies lesser-studied proteins with strong ESR1 and EZH2 associations, offering new leads for breast cancer drug discovery and precision medicine. These insights can help refine current clinical decision frameworks and guide the development of more personalized, precision-targeted therapies for breast cancer patients who currently lack clear genomic-based guidance.

## Methods

### Data Retrieval

Mutation data was obtained from DepMap v24Q4(17,18) and TCGA v2(19). Gene expression (RNA sequencing) and CRISPR-mediated knockdown data were obtained from DepMap. CRISPR data was obtained as a gene dependency score, where higher scores indicate increased dependency on ESR1/EZH2. Drug sensitivity data was obtained from the Cancer Therapeutics Research Portal (CTRP)(21–23).

### Classification of Rare vs Frequent mutations

Clinical mutation data was retrieved from OncoKB(24) and ClinVar(25). Mutations were classified as “frequent” if they occurred in >8% of breast cancer cell lines(26), and as “known drug targets” if they had a targeted therapeutic with an OncoKB evidence level of 3A or higher (24).

### Gene Set Enrichment Analysis

Single-sample GSEA(27) was performed on targets of interest using mutation and gene expression data. The ESR1 pathway was defined using an experimentally curated MSigDB(28) gene set(29), containing >200 genes. The EZH2 pathway was defined using an experimentally curated MSigDB(28) gene set(30), containing >1000 genes. Samples were ranked as “ESR1/EZH2 High” if they fell within the top 15 percentile of GSEA scores for that dataset.

### Mapping to the Protein Data Bank and Alphafold

Variants were mapped to the Protein Data Bank per residue using the mmtf-Python workflow (31), and to AlphaFold v2 using a previously developed workflow(32). Structural information was available for 17,852 variants across 8,886 genes for 69 breast cancer cell lines from DepMap, and 38,797 variants across 12,696 genes for 965 tumors in TCGA.

### ML Algorithm: Density-Based Clustering and Variant Classification

ML techniques were adapted from VAMOS(32). Features for ML included Cartesian coordinates and cluster information for each variant. We used a random one-third/two-thirds split for training and testing, respectively, with fivefold cross-validation. Training data was created by assigning mutations a preliminary classification of 0 (not associated) or 1 (associated) with a specific phenotype (e.g. top 15% GSEA score or 15% dependency score), and balanced via oversampling. Accuracy, recall, precision, F1 scores, AUC and MCC scores were calculated to assess performance (**SI 1-2**).

### Statistical Analysis of Clusters

The log odds that a cluster would contain a class “1” mutation was calculated using the formula:

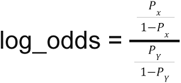

where *Px* is the probability of being a class “1” mutation and *Py* is the probability of being a class “0” mutation.

Priority scores were calculated by averaging log odds scores across all available data type combinations for each cluster.

### Identification of Statistically Significant Drug Associations

For each drug, cell lines containing class “1” clusters were compared to other cell lines, identifying cases where the mean AUC value was lower. Statistically significant differences (p<0.05) were determined using a Wilcoxon Rank Sum test, highlighting ML-cluster-associated drugs (**SI 3-20**).

For proteins with both class “1” clusters and other dense clusters, the Wilcoxon Rank Sum test was used to compare ML-cluster-associated drug sensitivity between dense-cluster-containing and non-dense-cluster-containing cell lines. For permutation analysis, n random mutations (where n = class “1” cluster size) were selected per protein, and the same test was applied. Permutations (100, 1,000, 10,000, and 100,000 iterations) were averaged per trial to assess statistical significance (**SI 3-20**).

### Code access

All code is available at github.com/Brunk-Lab/VAMOS_precision_oncology

## Results

### AI-Powered Annotation of Variants Linked to ESR1 and EZH2 Activity

Protein-coding variants are traditionally annotated based on impact on the function and structure of biomolecules(33). A variant is deemed pathogenic or benign depending on whether it alters the individual protein’s function(25). While this approach is useful for understanding molecular mechanisms, it falls short of capturing broader consequences of mutations on protein networks and cellular processes. For precision medicine, what truly matters is how these variants influence larger-scale biological functions, how they disrupt or reshape cellular pathways and, ultimately, how they determine an individual’s response to therapy. To bridge this gap, we need a more systematic approach for assessing functional consequences of variants beyond single-protein effects.

A comprehensive assessment of genome-wide variant effects requires large-scale data integration across multiple levels of biological organization. Fortunately, we are in an era of unprecedented data availability, with resources such as DepMap(17,18) and TCGA(19) offering extensive multi-omic datasets for functional variant interpretation. DepMap provides molecular profiles for >2,000 cancer cell lines, encompassing whole-genome sequencing (WGS), transcriptomics, proteomics, metabolomics, CRISPR-knockdown screens, and drug sensitivity data. Meanwhile, TCGA offers genomic, transcriptomic, and proteomic data across >11,000 tumor samples, with additional clinical metadata linking mutations to patient outcomes. Beyond these datasets, advances in protein structure modeling, particularly AlphaFold(20), now allow us to map thousands of variants directly onto three-dimensional protein structures, revealing insights into structure-function relationships. By identifying how mutations cluster within specific structural regions across tumors, we can move beyond simple frequency-based annotations, and instead, group mutations based on structural and functional context, a highly effective strategy for predicting broader impact on cellular processes(32).

To systematically link variants to the pathways they most likely influence, we developed VAMOS (Variant Annotation through Multi-Omic Signatures)(32), an ML-based framework that uncovers mutation-driven effects on cellular networks (**Fig. 1a**). Using mutation and transcriptional data from DepMap and TCGA, we trained an AI/ML model to recognize patterns in how mutations cluster in 3D protein space and to determine which clusters correlate most strongly with specific transcriptional and signaling activities. This approach demonstrated >90% accuracy in predicting pathway-level transcriptional outcomes of key transcription factors: NRF2 and MYC.

**Figure 1.**
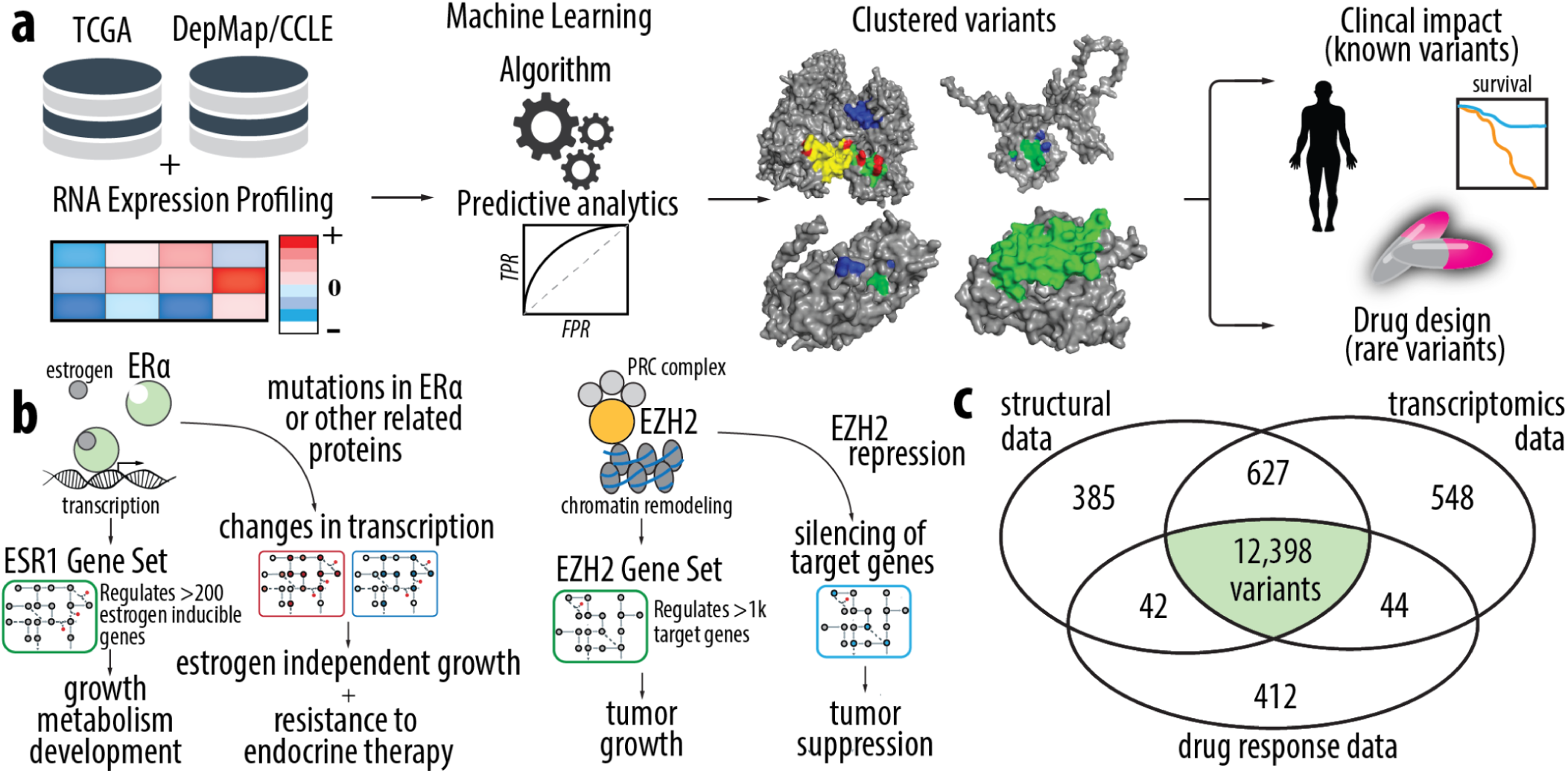
a) Overview of the ML pipeline to identify clinically relevant variants. Predictive analytics are performed on the algorithm to determine the quality of predictions. The output of the algorithm is clinically relevant dense variant clusters. These clusters are used to prioritize treatment options for known frequent variants, and identify novel treatment options for rare and understudied variants. b) This study looks at target pathway activity for ESR1 (encoding the ER) and EZH2. ESR1 has over 200 known target genes. ER is a steroid hormone that binds chromatin to regulate gene expression in response to estrogen. ER-activity is a primary determinant of intrinsic breast cancer biology, measured by gene expression profiles and clinical outcomes (recurrence and survival), and ER-inhibiting drugs are first line therapy for ER-positive tumors(34). EZH2 has over 1k known target genes, and is involved in epigenetic regulation primarily via chromatin remodeling, making it an exciting novel target for drug discovery(35). Mutations in these proteins (or other proteins) can lead to alterations in these pathways, and ultimately resistance to chemotherapeutics and hormone therapies. c) Variants from DepMap(17,18) with protein structural information, transcriptional information, and drug sensitivity information in the form of AUC values. 12,398 variants have all three types of information.

Here, we adapt this framework for translational science, focusing on key molecular drivers of breast cancer: ESR1 and EZH2. ESR1 (encoding estrogen receptor alpha) and EZH2 (enhancer of zeste homolog 2) are among the most important regulators of breast cancer progression and therapeutic response(34,35) (**Fig. 1b**). ESR1 mutations frequently emerge in endocrine-resistant tumors, driving estrogen-independent growth and limiting the efficacy of standard hormone therapies(34). EZH2, a histone methyltransferase, is a key epigenetic regulator implicated in tumor aggressiveness and drug resistance, particularly in luminal B and triple-negative breast cancers(35). Despite their importance, our ability to identify variants that alter the activity of these regulators remains limited. Identifying which variants influence ESR1 and EZH2 activity, and how they influence drug responses, would significantly enhance precision medicine strategies by enabling targeted therapeutic interventions.

Using our AI/ML framework, we systematically annotate >12,000 variants across 1034 breast cancer genomes, integrating RNA expression, single nucleotide variant (SNV) data, and drug response profiles (**Fig. 1c**). This multimodal approach allows us to rapidly categorize variants not only based on their structural and functional properties but also on their downstream transcriptional and therapeutic implications. This approach identifies both known and uncharacterized functionally relevant mutations that strongly associate with ESR1 or EZH2 activity. Furthermore, our framework reveals how these mutations influence drug sensitivity, opening new avenues for biomarker-driven treatment strategies in breast cancer.

### From Unknown to Actionable: AI/ML for Rapid Annotation of Rare Cancer Variants

Cancer mutations follow a long-tail distribution(36), where a small subset of mutations occurs frequently across the population and is well-characterized, while the vast majority are rare and largely unstudied (**Fig. 2a**). The current paradigm for variant annotation has prioritized frequent mutations, leading to major breakthroughs in targeted therapies that benefit hundreds of thousands of patients. However, 65% of breast cancer patients still lack frontline targeted care because they do not carry “actionable” mutations, or mutations with known functional impacts that can be directly linked to available therapies(24). We hypothesize that by shifting the focus from mutation frequency to structural context in 3D protein space, we can expand the set of functionally annotated rare variants, offering new opportunities for repurposing existing therapies and improving precision oncology.

**Figure 2.**
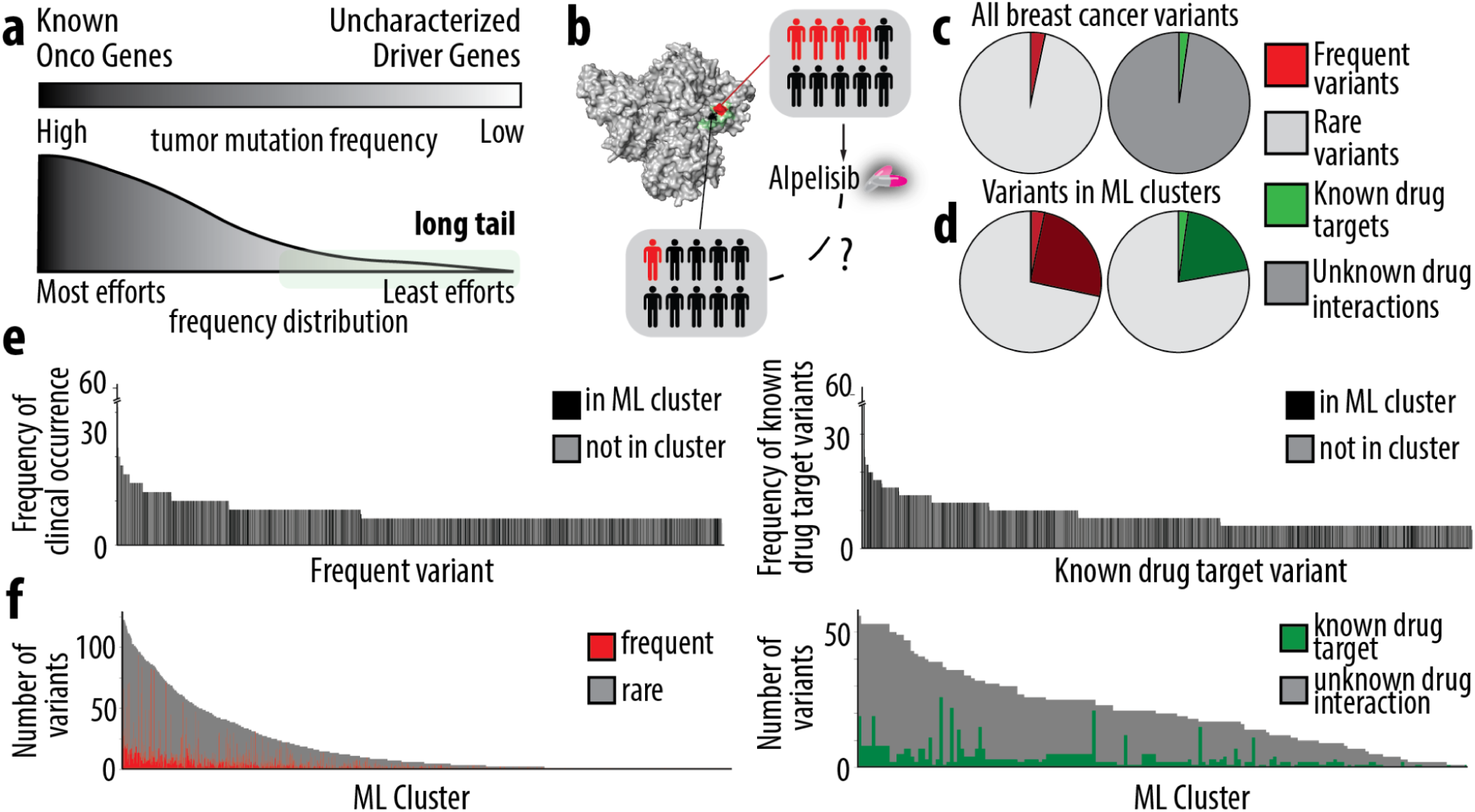
a) The “long tail” of distribution in cancer variants refers to a well known pattern in which a small number of genetic mutations are highly recurrent across patients, while a vast number of mutations occur infrequently. b) Frequent and rare variants may occur in the same dense cluster on a protein. Frequent variants are well studied and may have drugs that target them specifically. It is not yet known whether rare variants in proximity of frequent variants have the same biological and clinical relevance. c) The percentage of variants that are frequently occurring (left) or known drug targets (right) across all breast cancer cell lines and tumors. d) The percentage of variants that are frequently occurring or in the same cluster as a frequently occurring variant (left) or known drug targets or in the same cluster and a known drug target variant (right) across all breast cancer cell lines and tumors. e) The frequency of clinical occurrence for all variants in OncoKB(24) and ClinVar(25), variants present in ML identified dense clusters are highlighted in black (left). The frequency of known drug target variants for all variants in OncoKB(24) and ClinVar(25), variants present in ML identified dense clusters are highlighted in black (right). f) The number of frequently occurring variants in each ML identified dense clusters (left). The number of known drug target variants in each ML identified dense cluster (right).

To illustrate this approach, we use PIK3CA, a key oncogene frequently mutated in ER+ breast cancer. Well-known hotspot mutations, such as H1047R and E545K, are common across tumors and have well-established targeted therapies(37) (e.g., PI3K inhibitors). These therapies have shown significant clinical benefit for patients with these mutations, guiding treatment decisions in 15-20% of breast cancer cases(38,39). However, within the same structural region of the PIK3CA protein, we identified previously uncharacterized rare variants that are not currently considered actionable (**Fig. 2b**). Our goal is to use our AI/ML framework to determine whether rare variants exhibit similar structural and functional behavior to well-characterized mutations. If these rare mutations cluster with known pathogenic variants in 3D protein space and share functional impact, patients harboring them could potentially benefit from existing targeted therapies, expanding the reach of precision medicine to a broader patient population.

To estimate the prevalence of rare variants in breast cancer genomes, we analyzed 69 breast cancer cell lines and 965 tumor samples from DepMap(17,18) and TCGA(19) datasets, and mapped them to variant annotation databases, such as OncoKB(24), and ClinVar(25). Within this dataset, we identified 732 frequent variants, accounting for 5.06% of total mutations observed in this breast cancer subset (**Fig. 2c)**. By incorporating spatial clustering within 3D protein structures, we reduced the number of functionally unannotated variants by 35.6%, significantly enhancing our ability to infer their biological relevance. If these rare variants exhibit functional similarities to well-established mutations within their structural clusters, this approach could accelerate functional annotation without needing direct experimental validation. Furthermore, this approach reduced the number of variants not linked to targeted therapies by 23.4% (**Fig. 2d**), offering a clear opportunity for repurposing existing therapies to treat tumors harboring these previously unclassified mutations.

Interestingly, when mapping mutations onto the long-tail distribution of breast cancer genomes, we observed that rare variants were broadly distributed rather than clustering with frequent mutations or variants with targeted therapies, highlighting that our approach provides an independent layer of information beyond frequency-based classification (**Fig. 2e**). We assessed the composition of each AI/ML-identified cluster, examining the ratio of frequent vs. rare variants (**Fig. 2f**). The majority of mutation clusters contained a mix of well-characterized and understudied variants. This same principle applied to variants with available targeted therapies, suggesting that patients harboring rare variants in these clusters may respond to the same therapies as those with well-established, clinically actionable mutations.

### PIK3CA Variant Clusters as Predictors of ESR1 Activity and Therapy Response

To systematically identify variant clusters associated with ESR1 and EZH2 transcriptional activity, we quantified ESR1 activity using the collective expression of >200 ESR1 target genes, which is strongly correlated with sensitivity to endocrine therapy, across 69 breast cancer cell lines and 965 breast cancer tumor samples. We applied gene set enrichment analysis(27) (GSEA) to compute a standardized ESR1 activity score, which served as a proxy for ESR1 transcriptional activity(29). Using our AI/ML framework, we identified variant clusters that were most strongly correlated with ESR1 activity, rank-ordering proteins based on their log-odds of containing a mutation associated with ESR1 upregulation. We applied the same methodology to identify variant clusters linked to EZH2 activity, allowing us to systematically uncover mutations that associate with the activities of these key transcriptional regulators (**Methods**).

PIK3CA encodes the catalytic subunit of phosphatidylinositol 3-kinase (PI3K), a key regulator of cell growth, proliferation, and survival. Activating PIK3CA mutations enhance PI3K/AKT signaling, promoting tumor progression and therapy resistance(40). Our analysis identified several distinct PIK3CA variant clusters with high log-odds of association with ESR1 activity (**Fig. 3a**), including: (1) Cluster 1: Aligned with known ESR1-associated variants; (2) Cluster 2: Associated with upregulated ESR1 activity, with several previously annotated pathogenic variants; (3) Cluster 3: Linked to EZH2 upregulation, with no known variants of significance. Among these, Cluster 2 contained E545K, a well-characterized gain-of-function mutation(41), with 7.5% frequency across breast cancer genomes. The E545K mutation is known to enhance PI3K signaling, leading to constitutive activation(42) and persistent AKT activation, which drives tumorigenesis and therapy resistance(43). Because receptor subtypes guide treatment algorithms, we looked at subtype distribution by cluster. Cluster 1 is enriched for ER+/luminal which was expected given the identification using ER-activity signatures, while cluster 2 was more heterogenous, demonstrating the potential of these clusters as biomarkers of relevant driver signaling across subtypes (**SI 21**).

**Figure 3.**
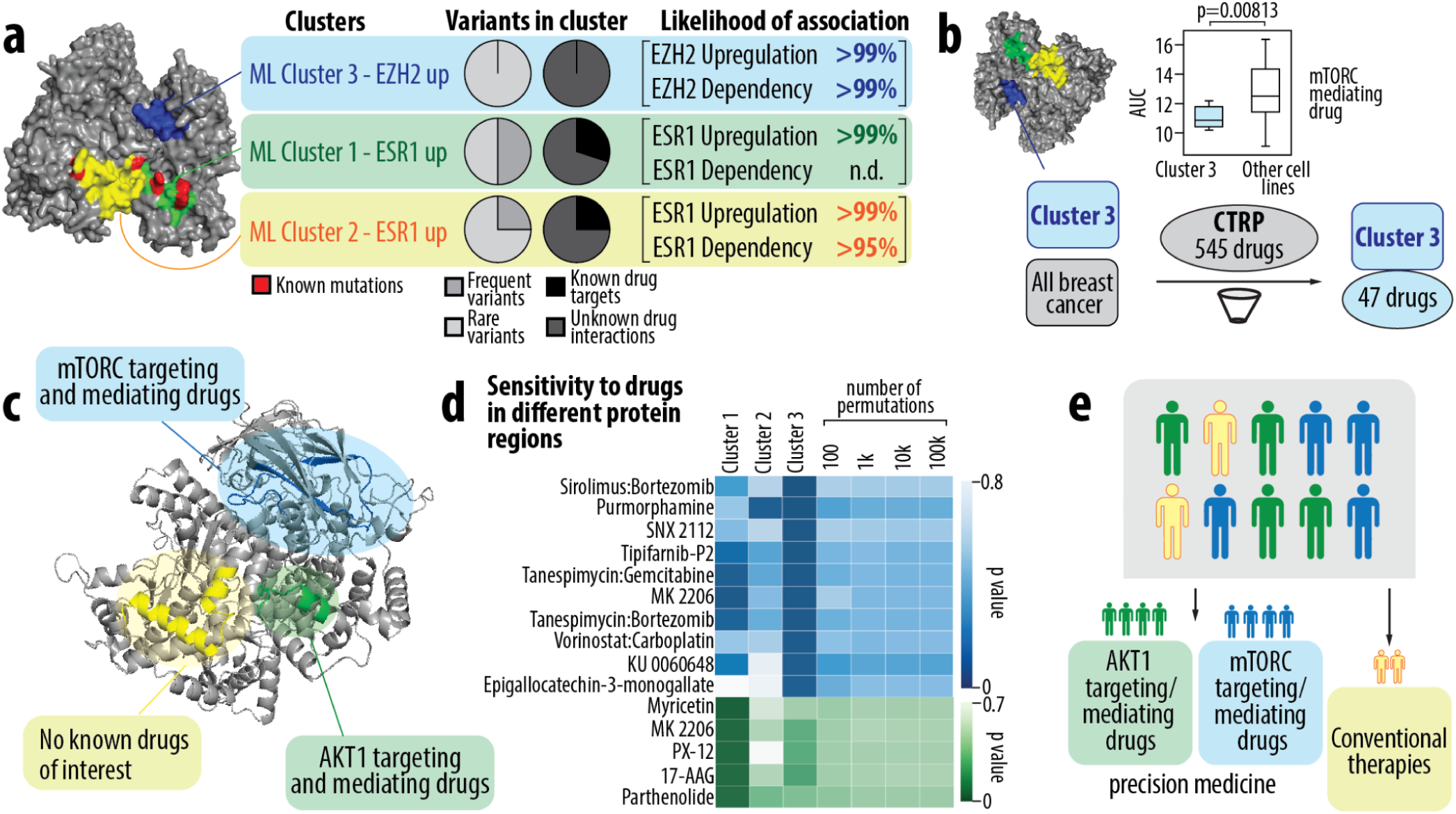
a) Three dense variant clusters in the PIK3CA protein are identified via our ML method. Cluster 1 (green) and Cluster 2 (yellow) variants are associated with upregulation of the ESR1 pathway. Cluster 3 (blue) variants are associated with upregulation of the EZH2 pathway. Variants with known clinical significance are highlighted in red. The center pie charts show the number of frequent vs rare variants and variants that are known drug targets vs variants with unknown drug interactions from Onco KB present in PIK3CA, and in each of the ML identified clusters. The likelihood that a cluster is associated with a given pathway is shown on the left. b) A schema showing the process for finding drugs with statistically different effects on cell lines containing the ML identified clusters, shown here for Cluster 3. Cell lines are sorted into two groups: those containing Cluster 3 variants, and those not containing Cluster 3 variants. Statistical analysis is performed to find all drugs that have lower AUC values for Cluster 3 cell lines and statistically different distributions compared to cell lines with no Cluster 3 mutations. c) Different regions of the protein are associated with increased sensitivity towards different drugs. Cell lines with Cluster 1 variants have greater sensitivity towards AKT1 targeting and mediating drugs. Cell lines with Cluster 2 variants do not have associations with any drugs. Cell lines with Cluster 3 variants have greater sensitivity towards mTORC targeting and mediating drugs. d) Heat maps showing the p-values generated by the statistical analysis described in part b for Cluster 1 and Cluster 3. Additional p values from random permutation analysis are also shown. e) These results can be used to inform precision medicine treatments for patients with mutations in the indicated clusters.

To further assess the functional significance of these variant clusters, we evaluated their impact on ESR1 and EZH2 pathway activity and cellular dependence on ESR1 or EZH2 using population-scale RNA sequencing data and CRISPR-mediated knockdown data of ESR1 and EZH2 data collected for 69 breast cancer cell lines (17,18). These analyses confirmed that variants within Clusters 1 and 2 were strongly predictive of ESR1 activity and ESR1 dependence in both cell lines and tumor samples (**SI 22-25**).

To assess the therapeutic implications of PIK3CA variant clusters, we integrated drug response profiles from the CTRP (Cancer Therapeutic Response Portal) database(21–23), which contains 545 drug response curves across 40 breast cancer cell lines. Using a Wilcoxon rank-sum test, we identified drugs that exhibited significantly different area under the curve (AUC) responses between samples harboring mutations in specific PIK3CA clusters and those without mutations in these clusters (**Fig. 3b**). The analysis revealed that cell lines with Cluster 3 variants displayed increased sensitivity to mTORC inhibitors and mTORC-mediating drugs (e.g. Sirolimus:Bortezomib, KU-0060648, Tipifarnib), while cell lines with Cluster 1 variants showed higher sensitivity to AKT inhibitors and AKT-modulating therapies (e.g. MK 2206, Myricetin, 17-AAG) (**Fig. 3c**). These findings suggest that the presence of specific PIK3CA variant clusters may dictate whether a tumor is more responsive to upstream (mTORC-targeting) or downstream (AKT-targeting) therapies (**Fig. 3d**).

Based on these findings, we propose a genomically-guided treatment approach in which patients with PIK3CA mutations are stratified according to these variant clusters (**Fig. 3e**). Patients with Cluster 1 mutations could benefit from AKT-targeting therapies, while those with Cluster 3 mutations may respond better to mTORC-based treatments. Applying this framework across 1034 breast cancer patients, we estimate that 14.5% would be classified for mTORC-based therapy and 13% for AKT-targeted therapy.

### Divergent TP53 Variant Clusters Drive Opposing ESR1 Activity and Therapeutic Sensitivities in Breast Cancer

TP53 emerged as a high ranked regulator in our analysis, with distinct variant clusters correlating to opposite ESR1 activity patterns. Unlike PIK3CA, where 2/3 identified clusters were linked to an increase in ESR1 transcriptional activities, TP53 exhibited two functionally divergent clusters: one strongly associated with increased ESR1 activity (ESR1_up) and another linked to decreased ESR1 (ESR1_down) activity (**Fig. 4a**). This suggests that mutational positioning within the TP53 protein structure may point to fundamentally different phenotypic consequences, with likely distinct therapeutic implications in breast cancer patients (**SI 26**).

**Figure 4.**
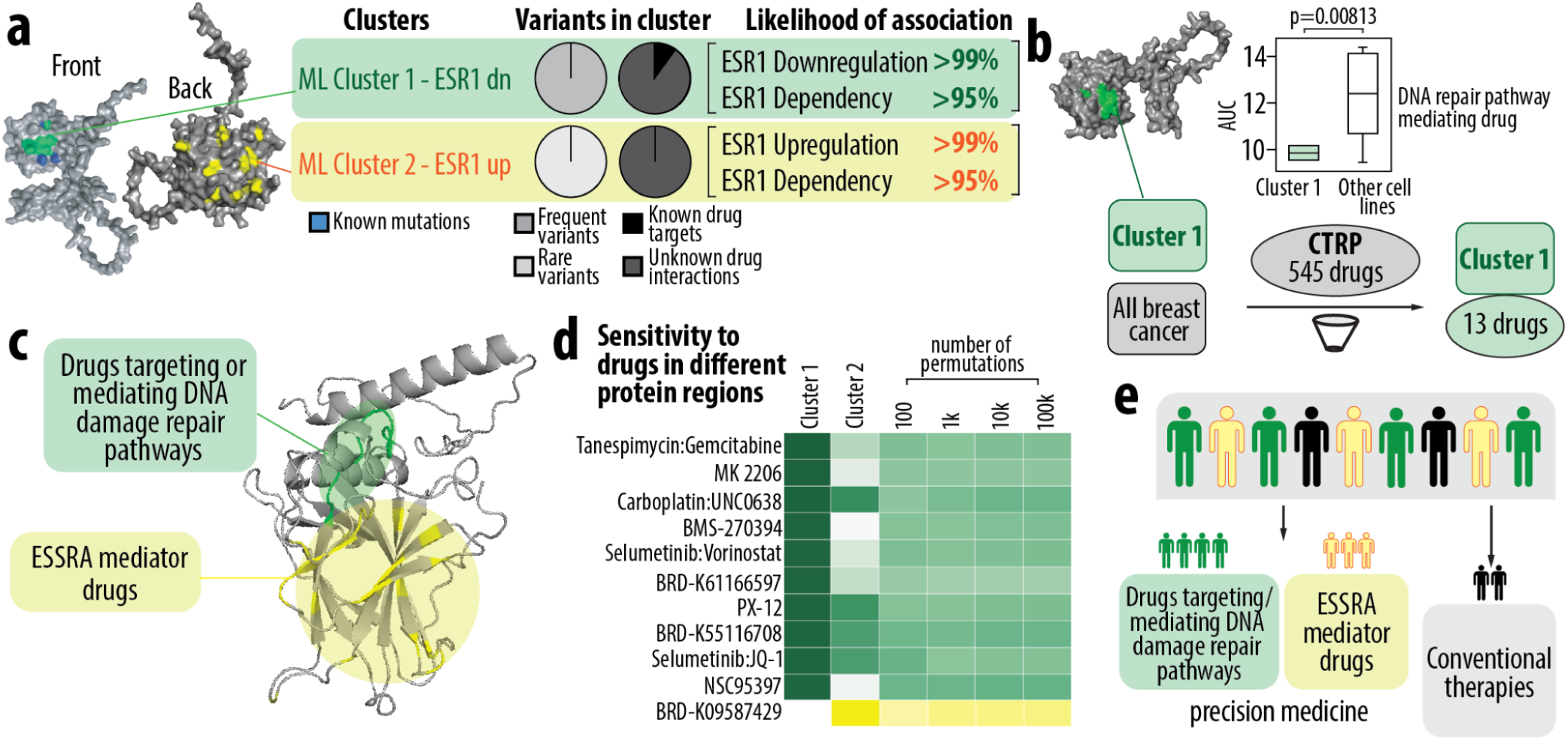
a) Two dense variant clusters in the TP53 protein are identified via our ML method. Cluster 1 (green) variants are associated with downregulation of the ESR1 pathway. Cluster 2 (yellow) variants are associated with upregulation of the ESR1 pathway. Variants with known clinical significance are highlighted in blue. The center pie charts show the number of frequent vs rare variants and variants that are known drug targets vs variants with unknown drug interactions from Onco KB present in TP53, and in each of the ML identified clusters. The likelihood that a cluster is associated with a given pathway is shown on the left. b) A schema showing the process for finding drugs with statistically different effects on cell lines containing the ML identified clusters, shown here for Cluster 1. Cell lines are sorted into two groups: those containing Cluster 1 variants, and those not containing Cluster 1 variants. Statistical analysis is performed to find all drugs that have lower AUC values for Cluster 1 cell lines and statistically different distributions compared to cell lines with no Cluster 1 mutations. c) Different regions of the protein are associated with increased sensitivity towards different drugs. Cell lines with Cluster 1 variants have greater sensitivity towards drugs targeting or mediating DNA damage repair pathways. Cell lines with Cluster 2 variants have greater sensitivity towards ESSRA mediator drugs. d) Heat maps showing the p-values generated by the statistical analysis described in part b for Cluster 1 and Cluster 2. Additional p values from random permutation analysis are also shown. e) These results can be used to inform precision medicine treatments for patients with mutations in the indicated clusters.

Both Cluster 1 and Cluster 2 were enriched for frequently occurring TP53 mutations, yet only a small fraction of these mutations are associated with targeted therapies (**Fig. 4a**). To determine whether variants within TP53 clusters represent loss-of-function (LOF) or gain-of-function (GOF) mutations, we integrated data from multiplex assays of variant effects (MAVE)(44,45), biochemical databases (BRENDA)(46), and oncogenic annotation platforms such as ClinVar(25) and OncoKB(24). Our analysis revealed that Cluster 1 (ESR1_down), was predominantly composed of LOF mutations (**SI 28**). Loss of TP53 function is known to compromise DNA damage response pathways, leading to uncontrolled proliferation and resistance to apoptosis(47). This loss of function has significant therapeutic implications, particularly for DNA-damaging agents and epigenetic modulators.

We examined the distribution of these variant clusters across breast cancer subtypes. As expected, Cluster 1 (ESR1_down) was predominantly found in basal-subtype tumors, while Cluster 2 (ESR1_up) was enriched in luminal-subtype tumors. However, ER status alone did not strongly stratify these clusters, likely because the classification captures ER activity rather than ER expression. These findings suggest that TP53 mutations in different structural pockets are associated with distinct molecular subtypes, a pattern that is not fully captured by traditional receptor-based classification. This highlights a layer of genomic heterogeneity that may influence tumor biology and treatment responses beyond what is accounted for by receptor status alone.

To validate the functional impact of these variant clusters, we analyzed the log-odds of their association with ESR1_up, ESR1_down, and ESR1 dependency. In both breast cancer cell lines and tumor samples, these TP53 clusters strongly correlated with their respective ESR1 phenotypes, reinforcing the biological relevance of their structural clustering and ranking (**SI 29**). We looked for differential drug sensitivity profiles for each cluster; we found 13 drugs that differentially sensitized samples with variants in Cluster 1 (**Fig. 4b**) and 1 drug for Cluster 2.

Endocrine therapies aim to block or degrade ESR1, reducing estrogen-driven tumor growth(48). However, in patients with TP53 mutations in Cluster 1, where ESR1 activity is downregulated, tumors may not be as dependent on estrogen signaling, making endocrine therapy less effective as a standalone treatment. Our findings suggest drugs targeting or mediating DNA damage repair pathways show significantly higher sensitizing effects on cell lines with variants in Cluster 1 compared to Cluster 2 (**Fig. 4d**). For these TP53-mutant, ESR1_down samples, platinum-based chemotherapy (e.g., carboplatin, cisplatin) could be combined with EHMT1/2 inhibitors (UNC0638(49)) to impair DNA repair mechanisms further, leading to enhanced cytotoxicity and tumor cell death. Additionally, PARP inhibitors (e.g., olaparib, talazoparib) or ATR/CHK1 inhibitors (e.g., AZD6738, prexasertib) could be incorporated to exploit the DNA repair deficiencies in TP53-mutant, ESR1_down tumors.

Patients with Cluster 2 (ESR1_up) variants could receive endocrine therapy (AIs or SERDs) with CDK4/6 inhibitors, while those with Cluster 1 (ESR1_down) variants could be treated with a combination of platinum-based chemotherapy, DNA repair inhibitors, and epigenetic modulators. Such an approach could optimize treatment efficacy by aligning therapy with the tumor’s molecular vulnerabilities, providing a more personalized approach to breast cancer management (**Fig. 4e**).

### A Treasure Trove of Untapped Targets in Breast Cancer Precision Medicine

Beyond well-characterized cancer drivers such as PIK3CA and TP53 our AI/ML framework identified a number of lesser-known proteins (e.g. DST, MGAM, KMT2C, TUBA1A) that exhibit strong associations with ESR1 and EZH2 activity (**Fig. 5a)**, making them compelling candidates for further investigation into their biological function, druggability, and therapeutic potential.

**Figure 5.**
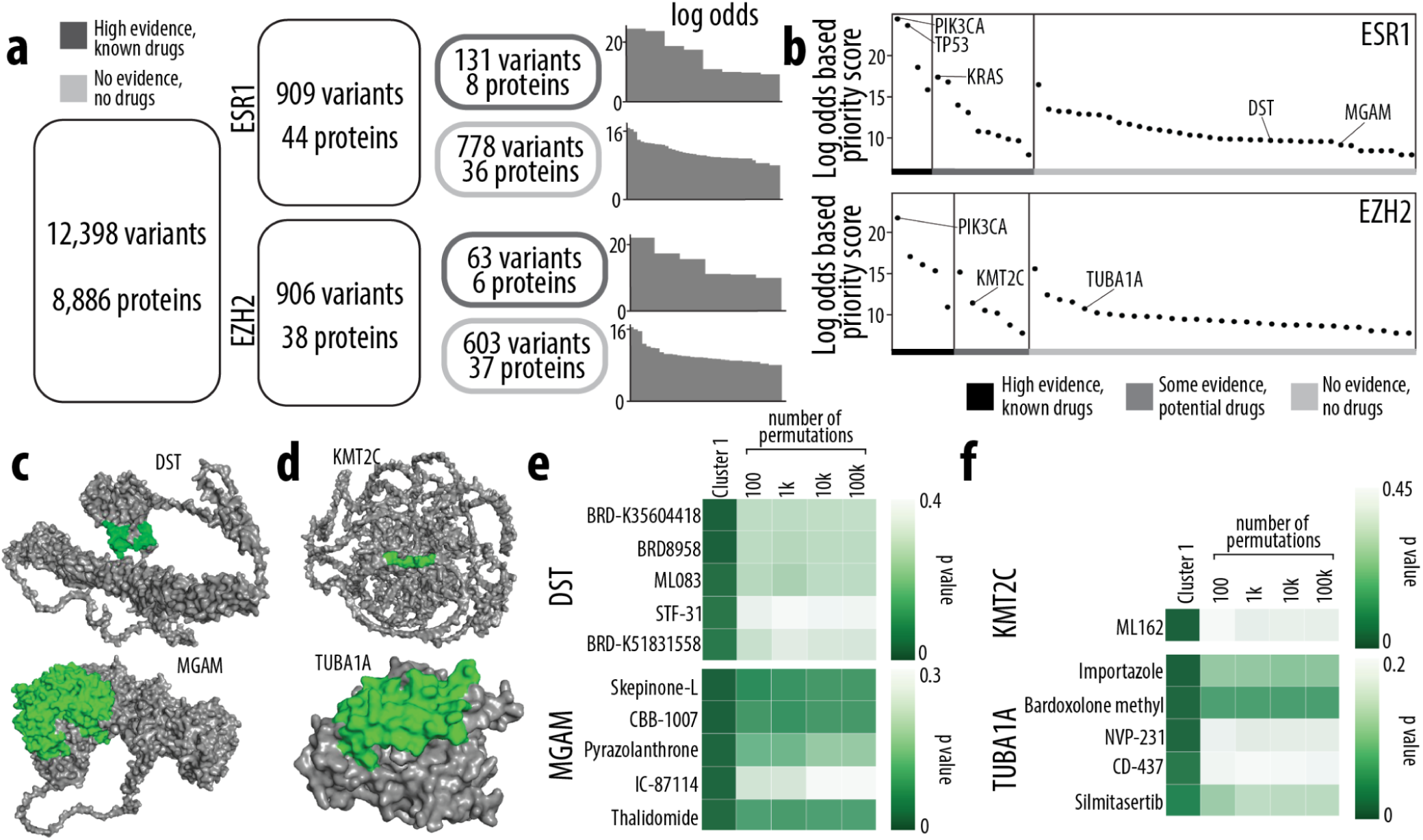
a) There are >12k variants across >8k proteins total with transcriptional data, protein structure data, and drug response data. Our workflow selected 909 variants across 44 proteins with a high likelihood of association with ESR activity, and 908 variants across 38 proteins with a high likelihood of association with EZH2 activity. The breakdown of known vs unknown variants and proteins for both ESR1 and EZH2 is shown. The logodds for each cluster identified by ML is shown on the right. b) This workflow can be used to prioritize new targets for study. Here, we show targets with a high logodds based priority score for future study. Targets are sorted into three bins. Bin 1 contains targets identified via our method that already have high evidence for impact in breast cancer, and already have known drugs. Bin 2 contains targets that have some evidence for impact in breast cancer, and have potential drugs. Bin 3 contains targets with no current evidence or drugs. c) Clusters in DST and MGAM, both Bin 3 targets, associated with ESR1 pathway changes. d) Clusters in KMT2C, a Bin 2 target, and TUBA1A, a Bin 3 target, associated with EZH2 pathway changes. e) Top 5 drugs identified using our method that have higher impact on variants in ML identified DST or MGAM clusters. f) Top drugs identified using our method that have higher impact on variants in ML identified KMT2C or TUBA1A clusters

Several of these proteins harbor rare and under-characterized mutations (887 variants across 51 proteins) that spatially cluster in 3D protein structures (**Fig. 5a**), mirroring the patterns seen in frequent, well-characterized oncogenes (e.g. PIK3CA and TP53). Despite their strong association with ESR1 and EZH2 transcriptional activities, many of these mutations have not been previously annotated as clinically actionable and remain absent from major oncogenic databases such as OncoKB(24) and ClinVar(25). However, our analysis suggests that these mutations may have functional consequences comparable to known driver mutations.

To further explore their clinical significance, we analyzed drug response profiles for these newly identified mutation clusters (**Fig. 5c**). Several ESR1-associated clusters were enriched for sensitivity to CDK4/6 inhibitors and PI3K inhibitors, while EZH2-associated clusters showed increased responsiveness to epigenetic modulators such as EZH2 inhibitors and BET inhibitors.

These findings emphasize the power of AI/ML in uncovering novel cancer drivers and highlight the need for further experimental validation to determine the precise molecular roles of these proteins in breast cancer pathogenesis. By systematically identifying untapped molecular targets with strong genomic, structural, and transcriptional relevance, our approach provides a roadmap for future investigations into their therapeutic potential, ultimately paving the way for more refined, biomarker-driven treatment strategies in breast cancer precision medicine.

## Discussion

Our AI/ML-driven framework represents a paradigm shift in how we classify and interpret genomic variants in breast cancer. By moving beyond frequency-based annotations to a structure-informed, functional approach, we identify previously unrecognized variants that strongly associate with ESR1 and EZH2 activity, highlighting new potential biomarkers and therapeutic targets. This approach expands the number of actionable mutations, offering new opportunities for patient stratification, drug repurposing, and precision-guided therapy selection.

Clinically, this work provides a scalable method to rapidly assess the functional impact of thousands of mutations, addressing a major limitation in current variant annotation strategies, which are slow and often limited to common mutations. Our findings demonstrate that rare or understudied mutations can cluster in the same regions of proteins as key oncogenic variants, emphasizing the need to reconsider how patients are classified for treatment. By integrating transcriptional, structural, and drug response data, we offer a framework to better predict which patients might benefit from endocrine therapy, PI3K/AKT inhibitors, or epigenetic modulators.

Moving forward, clinical validation of these variant clusters will be essential to confirm their relevance in enhancing treatment outcomes. Incorporating patient outcome data will further refine predictions and help identify which mutations contribute to therapy resistance or sensitivity. Additionally, this framework can be expanded beyond breast cancer, offering a generalizable AI-driven strategy for interpreting variants across multiple cancer types.

## Acknowledgements

The authors thank the UNC Longleaf Computer Cluster and Informatics team for their support in code optimization. This work was supported by funding from the Computational Medicine Pilot Award (no reference number) and the PhRMA Foundation Drug Discovery Faculty Starter Grant (no reference number).

## Author Contributions

Conceptualization, E.B. ; methodology, E.B., K.S. ; formal analysis, K.S., Y.W., P.S., E.B.; funding acquisition, E.B.; investigation, K.S., P.S., E.B.; resources, E.B.; supervision, E.B.; validation, K.S.; visualization, K.S., P.S., E.B.; writing—original draft, E.B., K.S., P.S.; writing—review and editing, All authors. All authors have read and agreed to the published version of the manuscript.

