## Supplementary material for "AI-Driven Variant Annotation for Precision Oncology in Breast Cancer": SI

Proteins, Genomics, Proteomics, Therapeutics, Algorithms, Classification, Computational Biology, Cancers

**SUPPORTING INFORMATION**

**SI 1:** Machine learning scores for the model. Scores are averages from 5-fold cross validation

| **Test Scores** | **Cell lines + RNAseq - ESR1 upregulated pathway** | **Cell lines + RNAseq - ESR1 downregulated pathway** | **Tumors + RNAseq- ESR1 upregulated pathway** | **Tumors + RNAseq- ESR1 downregulated pathway** | **Cell lines + CRISPR - ESR1 dependency** |
|---|---|---|---|---|---|
| **Accuracy** | 0.90696 | 0.85567 | 0.90317 | 0.82167 | 0.99146 |
| **Precision** | 0.84972 | 0.88673 | 0.83575 | 0.84869 | 0.98377 |
| **Recall** | 0.99711 | 0.97628 | 0.99729 | 0.99441 | 0.99971 |
| **F1** | 0.91752 | 0.80630 | 0.90939 | 0.85421 | 0.99167 |
| **MCC** | 0.82615 | 0.86082 | 0.82177 | 0.87956 | 0.98305 |

| Train Scores | Cell lines + RNAseq - ESR1 upregulated pathway | Cell lines + RNAseq - ESR1 downregulated pathway | Tumors + RNAseq- ESR1 upregulated pathway | Tumors + RNAseq- ESR1 downregulated pathway | Cell lines + CRISPR - ESR1 dependency |
|---|---|---|---|---|---|
| **Accuracy** | 0.98002 | 0.96702 | 0.98018 | 0.97096 | 0.99722 |
| **Precision** | 0.96526 | 0.95390 | 0.96321 | 0.95240 | 0.99484 |
| **Recall** | 0.99739 | 0.98424 | 0.99740 | 0.99441 | 0.99971 |
| **F1** | 0.98106 | 0.96883 | 0.98001 | 0.97295 | 0.99727 |
| **MCC** | 0.96051 | 0.93435 | 0.96095 | 0.94265 | 0.99444 |

| Test Scores | Cell lines + RNAseq - EZH2 upregulated pathway | Cell lines + RNAseq - EZH2 downregulated pathway | Tumors + RNAseq- EZH2 upregulated pathway | Tumors + RNAseq- EZH2 downregulated pathway | Cell lines + CRISPR - EZH2 dependency |
|---|---|---|---|---|---|
| **Accuracy** | 0.86267 | 0.97169 | 0.97069 | 0.82502 | 0.91480 |
| **Precision** | 0.89088 | 0.94694 | 0.94590 | 0.85049 | 0.85531 |
| **Recall** | 0.99554 | 0.99935 | 0.99850 | 0.99458 | 0.99769 |
| **F1** | 0.88147 | 0.97242 | 0.97148 | 0.85546 | 0.92102 |
| **MCC** | 0.85064 | 0.94486 | 0.94285 | 0.88642 | 0.84137 |

| Train Scores | Cell lines + RNAseq - EZH2 upregulated pathway | Cell lines + RNAseq - EZH2 downregulated pathway | Tumors + RNAseq- EZH2 upregulated pathway | Tumors + RNAseq- EZH2 downregulated pathway | Cell lines + CRISPR - EZH2 dependency |
|---|---|---|---|---|---|
| **Accuracy** | 0.97838 | 0.99253 | 0.99160 | 0.97117 | 0.98054 |
| **Precision** | 0.96352 | 0.98587 | 0.98492 | 0.95216 | 0.96436 |
| **Recall** | 0.99554 | 0.99935 | 0.99849 | 0.99458 | 0.99779 |
| **F1** | 0.97927 | 0.99256 | 0.99166 | 0.97291 | 0.98079 |
| **MCC** | 0.95725 | 0.98514 | 0.98329 | 0.94315 | 0.96165 |

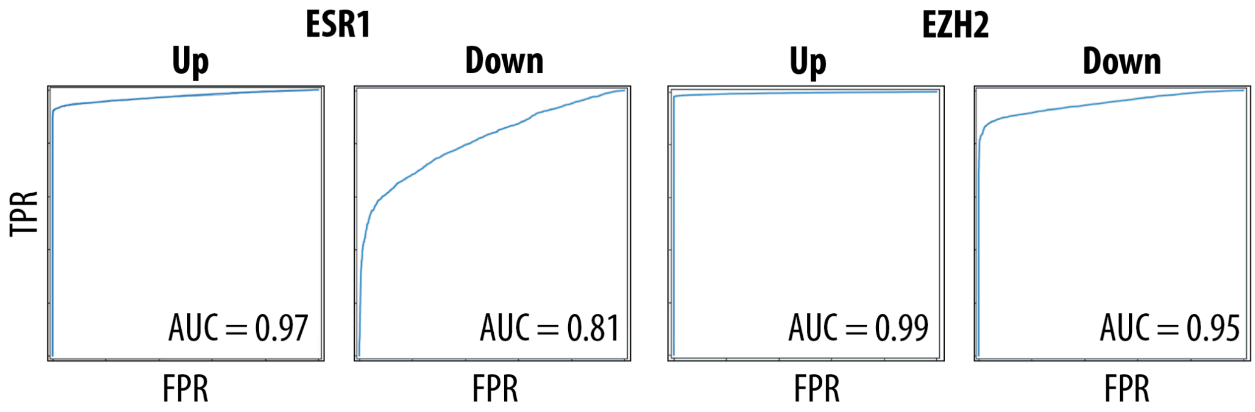


**SI 2:** Receiver operating curve (ROC) plots with Area under curve (AUC) scores for all combinations of machine learning performed

**SI 3:** Data for PIK3CA and the upregulated ESR1 pathway. P values for differences in AUC values for each drug between cell lines that contain variants in a given cluster and cell lines that don't.

| drug | VAMOS_clust | PIK3CA_other_1 | PIK3CA_other_2 |
|---|---|---|---|
| MYRICETIN | 0.01117 | 0.59951 | 0.45055 |
| BRD8899 | 0.02538 | 0.14702 | 0.78665 |
| SMER-3 | 0.02572 | 0.79400 | 0.97421 |
| MK 2206 | 0.02924 | 0.478521 | 0.27460 |
| PX-12 | 0.02944 | 0.753316 | 0.26289 |
| 17-AAG | 0.03448 | 0.485171 | 0.22667 |
| PARTHENOLIDE | 0.03917 | 0.326054 | 0.35555 |
| ML083 | 0.03917 | 0.160875 | 0.76828 |
| PRIMA-1 | 0.04706 | 0.552180 | 0.16830 |

**SI 4:** Data for PIK3CA and the upregulated ESR1 pathway. P values for differences in AUC values for each drug, where random clusters of the same size as the VAMOS cluster are chosen.

| drug | random_100_permutations | random_1000_permutations | random_10000_permutations | random_100000_permutations |
|---|---|---|---|---|
| ISOX:BORTEZOMIB | 0.46245 | 0.42125 | 0.43161 | 0.43597 |
| MYRICETIN | 0.42406 | 0.45764 | 0.45035 | 0.45377 |
| BRD8899 | 0.38642 | 0.39405 | 0.40177 | 0.40188 |
| SMER-3 | 0.50895 | 0.47250 | 0.46201 | 0.46167 |
| MK 2206 | 0.44039 | 0.47963 | 0.46764 | 0.46618 |
| PX-12 | 0.44410 | 0.45388 | 0.45618 | 0.45389 |
| 17-AAG | 0.44530 | 0.46194 | 0.46994 | 0.46834 |
| WAY-362450 | 0.38796 | 0.40597 | 0.38187 | 0.38551 |
| PARTHENOLIDE | 0.39097 | 0.43084 | 0.42937 | 0.43024 |
| ML083 | 0.52981 | 0.56523 | 0.55274 | 0.55232 |
| TANESPIMYCIN:GEMCITABINE | 0.39667 | 0.45499 | 0.45258 | 0.45101 |
| PRIMA-1 | 0.41666 | 0.40686 | 0.42417 | 0.41750 |

**SI 5:** Data for PIK3CA and the upregulated EZH2 pathway. P values for differences in AUC values for each drug between cell lines that contain variants in a given cluster and cell lines that don't.

| drug | PIK3CA_other_1 | PIK3CA_other_0 | VAMOS_clust |
|---|---|---|---|
| AY-22989 | 0.17806 | 0.95125 | 0.00451 |
| BRD-K63431240 | 0.52487 | 0.79621 | 0.00454 |
| BRD-K55116708 | 0.55655 | 0.59764 | 0.00826 |
| SIROLIMUS:BORTEZOMIB | 0.33100 | 0.60187 | 0.00981 |
| RG-108 | 0.59056 | 0.43800 | 0.01228 |
| MERCK60 | 0.36490 | 0.64576 | 0.01278 |
| BRD-A71883111 | 0.56283 | 0.00389 | 0.01324 |
| ML311 | 0.53635 | 0.22345 | 0.01378 |
| ISOX:BORTEZOMIB | 0.00702 | 0.17355 | 0.01569 |
| BRD-K07442505 | 0.84910 | 0.16804 | 0.01600 |
| DOCETAXEL:TANESPIMYCIN | 0.05507 | 0.45790 | 0.01842 |
| NAKITERPIOSIN | 0.92421 | 0.41021 | 0.01853 |
| PURMORPHAMINE | 0.52378 | 0.05132 | 0.01941 |
| NSC632839 | 0.73206 | 0.37591 | 0.01944 |
| SNX 2112 | 0.47555 | 0.61617 | 0.02141 |
| TIPIFARNIB-P2 | 0.10247 | 0.36503 | 0.02535 |
| BRD-A94377914 | 0.10247 | 0.65061 | 0.02535 |
| PRIMA-1-MET | 0.15304 | 0.95485 | 0.02535 |
| TUBASTATIN A | 0.15853 | 0.46685 | 0.02622 |
| TANESPIMYCIN:GEMCITABINE | 0.04653 | 0.38457 | 0.02686 |
| MK 2206 | 0.02924 | 0.47852 | 0.02746 |
| FGIN-1-27 | 0.17245 | 0.43668 | 0.02846 |
| BRD-K71935468 | 0.70711 | 0.58200 | 0.02867 |
| COMPOUND 1541A | 0.19491 | 0.40949 | 0.02924 |

| | | | |
|---|---|---|---|
| BRD-K49290616 | 0.10881 | 0.21704 | 0.02991 |
| BRD-K30019337 | 0.18145 | 0.31587 | 0.02991 |
| BRD-K78574327 | 0.18145 | 0.53709 | 0.02991 |
| BRD-K75293299 | 0.18145 | 0.31587 | 0.02991 |
| TANESPIMYCIN:BORTEZOMIB | 0.06464 | 0.40940 | 0.03301 |
| BRD-K70511574 | 0.53383 | 0.50425 | 0.03389 |
| VORINOSTAT:CARBOPLATIN | 0.52487 | 0.56079 | 0.03448 |
| ISOX | 0.36351 | 0.46490 | 0.03619 |
| DACARBAZINE | 0.31406 | 0.56816 | 0.03670 |
| MK-0683 | 0.95865 | 0.30255 | 0.03917 |
| BRD-K33199242 | 0.29048 | 0.34147 | 0.04216 |
| KU 0060648 | 0.17412 | 0.78996 | 0.04216 |
| BRD-K26531177 | 0.53909 | 0.08219 | 0.04296 |
| BRD-K28456706 | 0.29976 | 0.15730 | 0.04513 |
| BRD-K61166597 | 0.32457 | 0.96866 | 0.04513 |
| EPIGALLOCATECHIN-3-MONOGALLATE | 0.86333 | 0.80725 | 0.04776 |
| PL-DI | 0.05116 | 0.39015 | 0.04817 |
| WAY-362450 | 0.03904 | 0.16790 | 0.04831 |
| FLUOROURACIL | 0.72457 | 0.36113 | 0.04831 |
| BRD1812 | 0.87997 | 0.04787 | 0.04831 |
| BRD-K48334597 | 0.28505 | 1.00000 | 0.04840 |
| BRD-K96431673 | 0.28505 | 0.64343 | 0.04840 |
| TRIFLUOPERAZINE | 0.19962 | 0.65915 | 0.04980 |

**SI 6:** Data for PIK3CA and the upregulated EZH2 pathway. P values for differences in AUC values for each drug, where random clusters of the same size as the VAMOS cluster are chosen.

| drug | random_100_permutations | random_1000_permutations | random_10000_permutations | random_100000_permutations |
|---|---|---|---|---|
| AY-22989 | 0.31940 | 0.32524 | 0.32607 | 0.32343 |
| BRD-K63431240 | 0.46905 | 0.42634 | 0.44050 | 0.43942 |
| BRD-K55116708 | 0.38979 | 0.40395 | 0.39577 | 0.39957 |
| SIROLIMUS:BORTEZOMIB | 0.57213 | 0.54997 | 0.55311 | 0.55221 |
| RG-108 | 0.33365 | 0.34504 | 0.36104 | 0.35584 |
| MERCK60 | 0.58276 | 0.52332 | 0.51834 | 0.51612 |
| BRD-A71883111 | 0.28381 | 0.28631 | 0.28654 | 0.28351 |
| ML311 | 0.61573 | 0.59797 | 0.58640 | 0.59286 |
| ISOX:BORTEZOMIB | 0.47718 | 0.42238 | 0.43495 | 0.43273 |
| BRD-K07442505 | 0.33540 | 0.32841 | 0.33912 | 0.34176 |
| DOCETAXEL:TANESPIMYCIN | 0.41715 | 0.43612 | 0.43754 | 0.43534 |
| NAKITERPIOSIN | 0.44197 | 0.39260 | 0.40537 | 0.40426 |
| PURMORPHAMINE | 0.35458 | 0.40681 | 0.41050 | 0.40808 |
| NSC632839 | 0.48561 | 0.45042 | 0.44012 | 0.44284 |
| SNX 2112 | 0.56553 | 0.54213 | 0.53187 | 0.53646 |
| TIPIFARNIB-P2 | 0.45117 | 0.47930 | 0.44393 | 0.44689 |
| BRD-A94377914 | 0.31739 | 0.28165 | 0.27886 | 0.28087 |
| PRIMA-1-MET | 0.50342 | 0.56273 | 0.56085 | 0.55915 |
| TUBASTATIN A | 0.32443 | 0.31319 | 0.31708 | 0.31785 |
| TANESPIMYCIN:GEMCITABINE | 0.42530 | 0.46495 | 0.44937 | 0.45117 |
| MK 2206 | 0.55374 | 0.46471 | 0.46391 | 0.46350 |
| FGIN-1-27 | 0.32189 | 0.29672 | 0.30153 | 0.30503 |
| BRD-K71935468 | 0.37382 | 0.35361 | 0.35705 | 0.35828 |
| COMPOUND 1541A | 0.39603 | 0.41858 | 0.43455 | 0.42933 |

| | | | | |
|---|---|---|---|---|
| BRD-K49290616 | 0.42036 | 0.43037 | 0.42843 | 0.42938 |
| BRD-K30019337 | 0.23848 | 0.26906 | 0.26907 | 0.27386 |
| BRD-K78574327 | 0.46309 | 0.40465 | 0.42572 | 0.42686 |
| BRD-K75293299 | 0.17917 | 0.28144 | 0.26253 | 0.26436 |
| TANESPIMYCIN:BORTEZOMIB | 0.42490 | 0.45832 | 0.44489 | 0.45066 |
| BRD-K70511574 | 0.46216 | 0.46710 | 0.47602 | 0.47862 |
| VORINOSTAT:CARBOPLATIN | 0.49867 | 0.46300 | 0.47967 | 0.47741 |
| ISOX | 0.50738 | 0.46766 | 0.47810 | 0.47735 |
| DACARBAZINE | 0.48529 | 0.48181 | 0.50419 | 0.50189 |
| MK-0683 | 0.39760 | 0.38570 | 0.39721 | 0.39798 |
| BRD-K33199242 | 0.49523 | 0.45984 | 0.45462 | 0.45270 |
| KU 0060648 | 0.32366 | 0.37022 | 0.36348 | 0.36138 |
| BRD-K26531177 | 0.31749 | 0.34800 | 0.32860 | 0.32735 |
| BRD-K28456706 | 0.40783 | 0.41024 | 0.40993 | 0.40860 |
| BRD-K61166597 | 0.43226 | 0.44191 | 0.43720 | 0.43944 |
| EPIGALLOCATECHIN-3-MONOGALLATE | 0.41096 | 0.48742 | 0.48621 | 0.48019 |
| PL-DI | 0.42858 | 0.42209 | 0.42302 | 0.42651 |
| WAY-362450 | 0.41253 | 0.38781 | 0.38624 | 0.38742 |
| FLUOROURACIL | 0.47306 | 0.47169 | 0.46426 | 0.46165 |
| BRD1812 | 0.39077 | 0.38626 | 0.40263 | 0.40014 |
| BRD-K48334597 | 0.35925 | 0.37791 | 0.38204 | 0.38014 |
| BRD-K96431673 | 0.41350 | 0.43670 | 0.42118 | 0.42095 |
| TRIFLUOPERAZINE | 0.41938 | 0.41795 | 0.41704 | 0.41944 |

**SI 7:** Data for TP53 and the downregulated ESR1 pathway. P values for differences in AUC values for each drug between cell lines that contain variants in a given cluster and cell lines that don't.

| drug | VAMOS_clust | TP53_other_1 | TP53_other_2 | TP53_other_3 | TP53_other_4 |
|---|---|---|---|---|---|
| TANESPIMYCIN:GEMCITABINE | 0.02585 | 0.32838 | 0.16128 | 0.59395 | 0.63526 |
| MK 2206 | 0.02993 | | 0.14780 | 0.65472 | 0.82306 |
| CARBOPLATIN:UNC0638 | 0.03379 | | 0.09229 | 0.28450 | 0.19175 |
| BMS-270394 | 0.03733 | 0.16006 | 0.20190 | 0.34897 | 0.95563 |
| SELUMETINIB:VORINOSTAT | 0.03931 | | 0.28642 | 0.76643 | 0.77621 |
| BRD-K61166597 | 0.03931 | | 0.56967 | 0.76643 | 0.67999 |
| PX-12 | 0.03931 | | 0.20083 | 0.48824 | 0.21592 |
| BRD-K55116708 | 0.04042 | | 0.21340 | 0.33288 | 0.27225 |
| MYRICETIN | 0.04164 | | 0.90989 | 0.79290 | 0.88006 |
| SELUMETINIB:JQ-1 | 0.04391 | 0.34897 | 0.15840 | 0.45370 | 0.26578 |
| BRD-K63431240 | 0.04391 | 0.34897 | 0.50180 | 0.85141 | 0.50433 |
| NSC95397 | 0.04556 | | 0.08268 | | 0.90989 |

**SI 8:** Data for TP53 and the downregulated ESR1 pathway. P values for differences in AUC values for each drug, where random clusters of the same size as the VAMOS cluster are chosen.

| drug | random_100_permutations | random_1000_permutations | random_10000_permutations | random_100000_permutations |
|---|---|---|---|---|
| TANESPIMYCIN:GEMCITABINE | 0.45695 | 0.48902 | 0.45951 | 0.45544 |
| MK 2206 | 0.49726 | 0.50468 | 0.49791 | 0.49965 |
| CARBOPLATIN:UNC0638 | 0.50088 | 0.44285 | 0.39742 | 0.39800 |
| BMS-270394 | 0.47148 | 0.48585 | 0.46222 | 0.45301 |
| SELUMETINIB:VORINOSTAT | 0.45148 | 0.46951 | 0.45476 | 0.44684 |
| BRD-K61166597 | 0.53035 | 0.57148 | 0.56640 | 0.56912 |
| PX-12 | 0.47503 | 0.47657 | 0.44779 | 0.44535 |
| BRD-K55116708 | 0.41318 | 0.42556 | 0.39652 | 0.40305 |
| MYRICETIN | 0.58145 | 0.55949 | 0.58161 | 0.58414 |
| SELUMETINIB:JQ-1 | 0.39087 | 0.49163 | 0.46547 | 0.46868 |
| BRD-K63431240 | 0.55545 | 0.56834 | 0.56445 | 0.56433 |
| NSC95397 | 0.40257 | 0.39914 | 0.37874 | 0.37929 |

**SI 9:** Data for TP53 and the upregulated ESR1 pathway. P values for differences in AUC values for each drug between cell lines that contain variants in a given cluster and cell lines that don't.

| drug | VAMOS_clust | TP53_other_1 | TP53_other_2 |
|---|---|---|---|
| BRD-K09587429 | 0.03887 | 0.23200 | 0.81107 |

**SI 10:** Data for TP53 and the upregulated ESR1 pathway. P values for differences in AUC values for each drug, where random clusters of the same size as the VAMOS cluster are chosen.

| drug | random_100_permutations | random_1000_permutations | random_10000_permutations | random_100000_permutations |
|---|---|---|---|---|
| BRD-K09587429 | 0.45415 | 0.33170 | 0.32429 | 0.31984 |

**SI 11:** Data for PTEN and the downregulated ESR1 pathway. P values for differences in AUC values for each drug between cell lines that contain variants in a given cluster and cell lines that don't.

| drug | VAMOS_clust | PTEN_other_1 |
|---|---|---|
| GSK525762A | 0.00443 | 0.53396 |
| VORINOSTAT:CARBOPLATIN | 0.00540 | 0.34897 |
| I-BET151 | 0.00737 | 0.92734 |
| DACARBAZINE | 0.00921 | 0.88519 |
| ML050 | 0.00921 | 0.22887 |
| BRD-K80183349 | 0.01128 | 0.92113 |
| BRD1812 | 0.01278 | 0.59650 |
| BMS-270394 | 0.01673 | 0.51208 |
| NECROSTATIN-7 | 0.01853 | 0.81966 |
| CARBOPLATIN:UNC0638 | 0.01941 | 0.20262 |
| ML029 | 0.02279 | 0.64646 |
| SELUMETINIB:VORINOSTAT | 0.02514 | 0.92113 |

**SI 12:** Data for PTEN and the downregulated ESR1 pathway. P values for differences in AUC values for each drug, where random clusters of the same size as the VAMOS cluster are chosen.

| drug | random_100_permutations | random_1000_permutations | random_10000_permutations | random_100000_permutations |
|---|---|---|---|---|
| GSK525762A | 0.00475 | 0.00475 | 0.01589 | 0.01594 |
| VORINOSTAT:CARBOPLATIN | 0.05041 | 0.05041 | 0.12835 | 0.12807 |
| I-BET151 | 0.01630 | 0.01630 | 0.04499 | 0.04466 |
| DACARBAZINE | 0.01799 | 0.01799 | 0.04702 | 0.04703 |
| ML050 | 0.09669 | 0.09669 | 0.20679 | 0.20604 |
| BRD-K80183349 | 0.02253 | 0.02253 | 0.05622 | 0.05590 |
| BRD1812 | 0.05577 | 0.05577 | 0.12045 | 0.12123 |
| BMS-270394 | 0.01446 | 0.01446 | 0.03823 | 0.03834 |
| NECROSTATIN-7 | 0.02867 | 0.02867 | 0.06394 | 0.06361 |
| CARBOPLATIN:UNC0638 | 0.16460 | 0.16460 | 0.29691 | 0.30139 |
| ML029 | 0.07774 | 0.07774 | 0.14796 | 0.14795 |
| SELUMETINIB:VORINOSTAT | 0.05507 | 0.05507 | 0.11283 | 0.11370 |

**SI 13:** Data for DST and the upregulated ESR1 pathway. P values for differences in AUC values for each drug between cell lines that contain variants in a given cluster and cell lines that don’t.

| drug | VAMOS_clust | DST_other_1 | DST_other_2 | DST_other_3 | DST_4 |
|---|---|---|---|---|---|
| BRD-K35604418 | 0.00813 | 0.13092 | 0.92920 | 0.30832 | 0.20287 |
| BRD8958 | 0.00953 | 0.11518 | 0.20758 | 0.76281 | 0.40081 |
| ML083 | 0.02578 | 0.11314 | 0.69206 | 0.32210 | 0.92113 |
| STF-31 | 0.03917 |  | 0.92113 | 0.42829 | 0.13749 |
| BRD-K51831558 | 0.04417 | 0.14413 | 0.85513 | 0.20124 |  |

**SI 14:** Data for DST and the upregulated ESR1 pathway. P values for differences in AUC values for each drug, where random clusters of the same size as the VAMOS cluster are chosen.

| drug | random_100_permutations | random_1000_permutations | random_10000_permutations | random_100000_permutations |
|---|---|---|---|---|
| BRD-K35604418 | 0.31663 | 0.31414 | 0.32111 | 0.32446 |
| BRD8958 | 0.31255 | 0.30140 | 0.31160 | 0.31356 |
| ML083 | 0.30521 | 0.28283 | 0.31485 | 0.31510 |
| STF-31 | 0.42477 | 0.45906 | 0.44369 | 0.44108 |
| BRD-K51831558 | 0.33714 | 0.39251 | 0.36973 | 0.36893 |

**SI 15:** Data for MGAM and the downregulated ESR1 pathway. P values for differences in AUC values for each drug between cell lines that contain variants in a given cluster and cell lines that don't.

| drug | VAMOS_clust | MGAM_other_1 | MGAM_other_2 | MGAM_other_3 |
|---|---|---|---|---|
| SKEPINONE-L | 0.02195 | 0.13167 | | 0.09725 |
| CBB-1007 | 0.02236 | 0.13429 | | 0.09792 |
| PYRAZOLANTHRONE | 0.02677 | 0.26829 | 0.08425 | 0.47032 |
| IC-87114 | 0.02677 | 0.13568 | 0.18946 | 0.36046 |
| THALIDOMIDE | 0.03104 | 0.18645 | | 0.09865 |
| BRD-K85133207 | 0.03322 | 0.10974 | 0.56662 | 0.09137 |
| SZ4TA2 | 0.03981 | 0.23714 | | 0.09792 |
| VX-680 | 0.04042 | 0.12134 | | 0.24528 |
| BRD-K90370028 | 0.04382 | 0.19337 | | 0.14046 |
| KU-60019 | 0.04407 | 0.09207 | 0.24038 | 0.59650 |
| KU 0060648 | 0.04831 | | 0.24038 | 0.09207 |
| CARBOPLATIN:UNC0638 | 0.04846 | | 0.14323 | 0.20262 |

**SI 16:** Data for MGAM and the downregulated ESR1 pathway. P values for differences in AUC values for each drug, where random clusters of the same size as the VAMOS cluster are chosen.

| drug | random_100_permutations | random_1000_permutations | random_10000_permutations | random_100000_permutations |
|---|---|---|---|---|
| SKEPINONE-L | 0.06988 | 0.08102 | 0.08931 | 0.08873 |
| CBB-1007 | 0.08722 | 0.08011 | 0.08946 | 0.08959 |
| PYRAZOLANTHRONE | 0.14035 | 0.14253 | 0.18026 | 0.17994 |
| IC-87114 | 0.24717 | 0.25179 | 0.31367 | 0.31449 |
| THALIDOMIDE | 0.10209 | 0.09558 | 0.10305 | 0.10393 |
| BRD-K85133207 | 0.26912 | 0.25663 | 0.30196 | 0.30177 |
| SZ4TA2 | 0.11463 | 0.10817 | 0.11691 | 0.11660 |
| VX-680 | 0.15994 | 0.15671 | 0.17262 | 0.17180 |
| BRD-K90370028 | 0.12303 | 0.12158 | 0.13104 | 0.13084 |
| KU-60019 | 0.33420 | 0.30522 | 0.36032 | 0.36182 |
| KU 0060648 | 0.18416 | 0.18919 | 0.23309 | 0.23357 |
| CARBOPLATIN:UNC0638 | 0.20723 | 0.22988 | 0.26997 | 0.27035 |

**SI 17:** Data for TUBA1A and the upregulated EZH2 pathway. P values for differences in AUC values for each drug between cell lines that contain variants in a given cluster and cell lines that don't.

| drug | VAMOS_clust | TUBA1A_other_1 | TUBA1A_other_2 |
|---|---|---|---|
| ERLOTINIB | 0.03917 | 0.00561 | 0.61886 |
| IMPORTAZOLE | 0.00672 | 0.00561 | 0.56967 |
| GEFITINIB | 0.06404 | 0.00573 | 0.30551 |
| PD 153035 | 0.13167 | 0.01198 | 0.18645 |
| BARDOXOLONE METHYL | 0.01128 | 0.01333 | 0.31977 |
| CANERTINIB | 0.03952 | 0.01436 | 0.42025 |
| CD-437 | 0.02377 | 0.01499 | 0.67860 |
| SILMITASERTIB | 0.03184 | 0.01499 | 0.30024 |
| BRD9876 | 0.04813 | 0.01917 | 0.16881 |
| CRA-032765 | 0.04024 | 0.02072 | 0.16994 |
| AFATINIB | 0.14071 | 0.02156 | 0.72234 |
| ISOEVODIAMINE | 0.14071 | 0.02156 | 0.83117 |
| FLUOROURACIL | 0.04831 | 0.02377 | 0.94494 |
| ML006 | 0.11549 | 0.02377 | 0.72986 |
| WZ4002 | 0.22780 | 0.02703 | 0.18645 |
| BRD-K24690302 | 0.03448 | 0.03001 | 0.94645 |
| NERATINIB | 0.04236 | 0.03250 | 0.64724 |
| NVP-231 | 0.01141 | 0.03501 | 0.48369 |
| SELUMETINIB:GDC-0941 | 0.06286 | 0.03670 | 0.18946 |

**SI 18:** Data for TUBA1A and the upregulated EZH2 pathway. P values for differences in AUC values for each drug, where random clusters of the same size as the VAMOS cluster are chosen.

| drug | random_100_permutations | random_1000_permutations | random_10000_permutations | random_100000_permutations |
|---|---|---|---|---|
| ERLOTINIB | 0.08405 | 0.09170 | 0.09312 | 0.09330 |
| IMPORTAZOLE | 0.11749 | 0.11602 | 0.10968 | 0.11124 |
| GEFITINIB | 0.07759 | 0.06843 | 0.07268 | 0.07232 |
| PD 153035 | 0.08220 | 0.07625 | 0.07579 | 0.07413 |
| BARDOXOLONE METHYL | 0.06384 | 0.06319 | 0.06168 | 0.06229 |
| CANERTINIB | 0.11791 | 0.13974 | 0.14140 | 0.14377 |
| CD-437 | 0.20451 | 0.21342 | 0.21126 | 0.20669 |
| SILMITASERTIB | 0.12376 | 0.14823 | 0.14510 | 0.14434 |
| BRD9876 | 0.16101 | 0.18574 | 0.18856 | 0.18626 |
| CRA-032765 | 0.16128 | 0.15272 | 0.15438 | 0.15550 |
| AFATINIB | 0.30914 | 0.27624 | 0.26710 | 0.27056 |
| ISOEVODIAMINE | 0.18412 | 0.22123 | 0.20946 | 0.21112 |
| FLUOROURACIL | 0.12854 | 0.16041 | 0.15379 | 0.15447 |
| ML006 | 0.13592 | 0.13091 | 0.13253 | 0.13166 |
| WZ4002 | 0.27731 | 0.26852 | 0.27168 | 0.26723 |
| BRD-K24690302 | 0.19759 | 0.20263 | 0.20250 | 0.20337 |
| NERATINIB | 0.21601 | 0.23588 | 0.23994 | 0.23637 |
| NVP-231 | 0.19899 | 0.18465 | 0.18648 | 0.18633 |
| SELUMETINIB:GDC-0941 | 0.11953 | 0.12073 | 0.12047 | 0.12029 |

**SI 19:** Data for KMT2C and the upregulated EZH2 pathway. P values for differences in AUC values for each drug between cell lines that contain variants in a given cluster and cell lines that don't.

| drug | VAMOS_clust | KMT2C_other_1 | KMT2C_other_2 | KMT2C_other_3 | KMT2C_other_4 |
|---|---|---|---|---|---|
| ML162 | 0.04391 | 0.94645 | 0.57415 | 0.68695 | |

**SI 20:** Data for KMT2C and the upregulated EZH2 pathway. P values for differences in AUC values for each drug, where random clusters of the same size as the VAMOS cluster are chosen.

| drug | random_100_permutations | random_1000_permutations | random_10000_permutations | random_100000_permutations |
|---|---|---|---|---|
| ML162 | 0.48898 | 0.43895 | 0.44687 | 0.44309 |

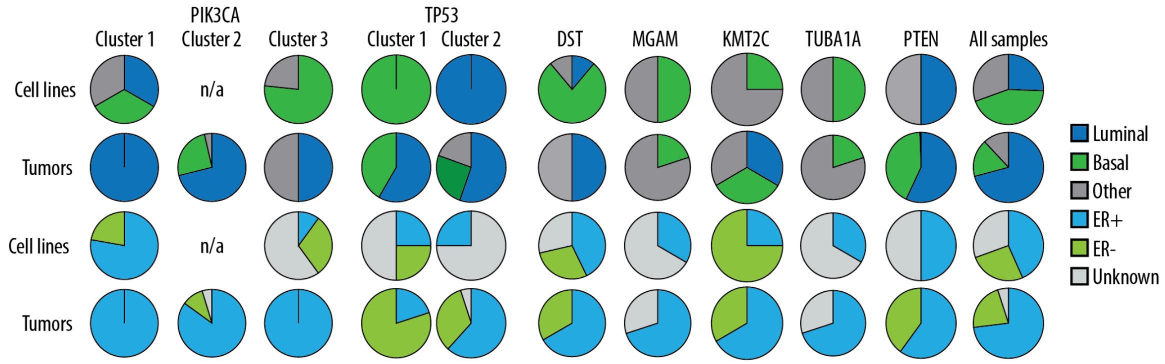


**SI 21:** Tissue type breakdown for all clusters

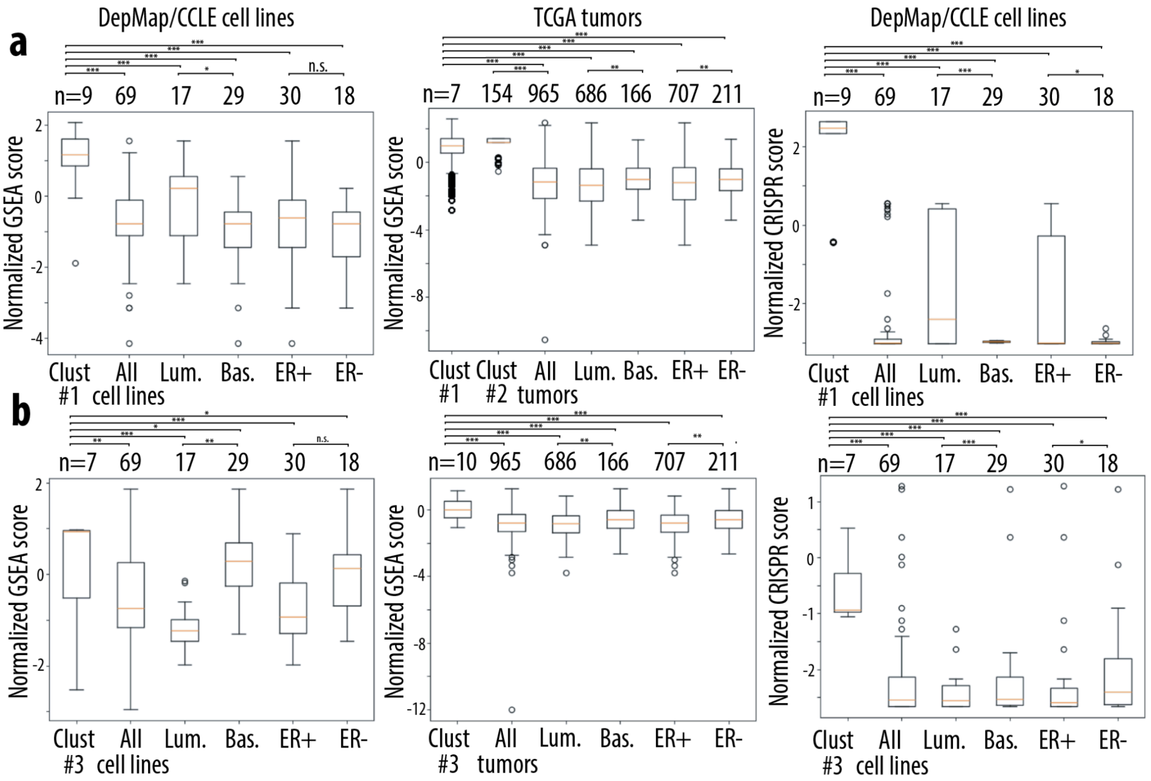


**SI 22:** a) Boxplots showing normalized GSEA (pathway expression) and normalized CRISPR dependency scores for cell lines and tumors with PIK3CA mutations in Machine Learning identified Cluster 1 and 2. These mutation clusters are associated with upregulated ESR1 activity. b) Boxplots showing normalized GSEA (pathway expression) and normalized CRISPR dependency scores for cell lines and tumors with

PIK3CA mutations in Machine Learning identified Cluster 3. These mutation clusters are associated with upregulated EZH2 activity.

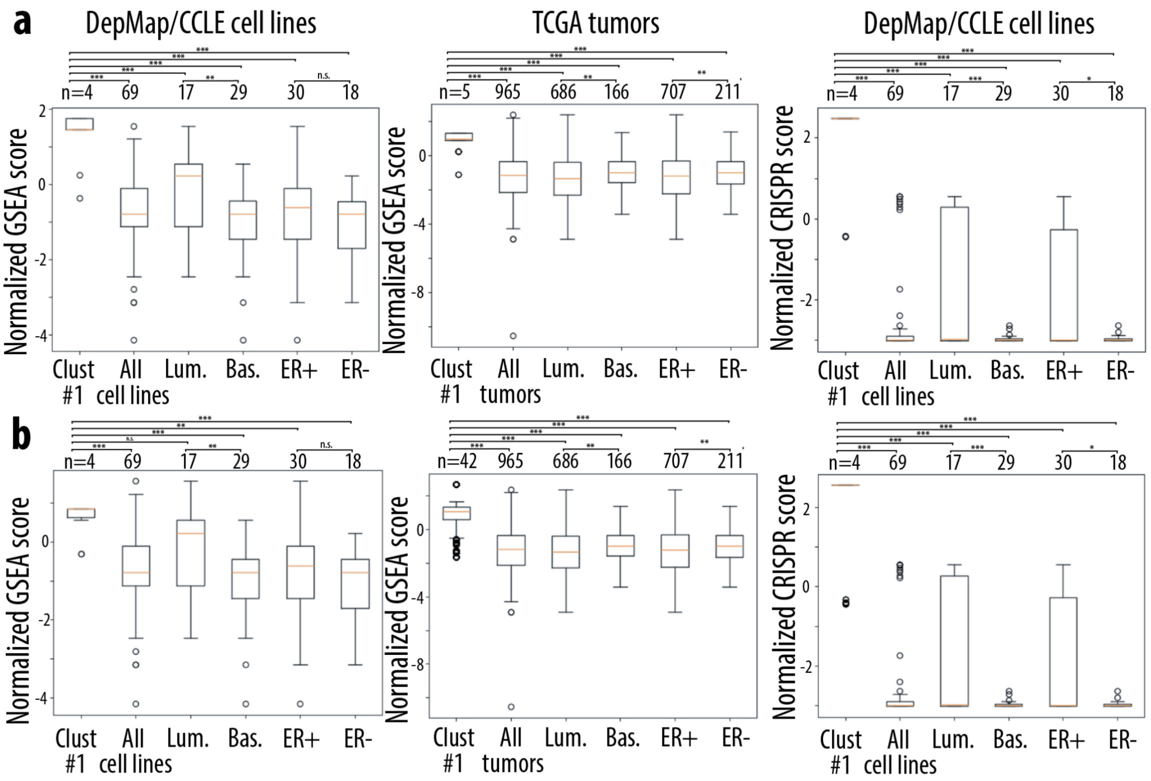


**SI 23:** a) Boxplots showing normalized GSEA (pathway expression) and normalized CRISPR dependency scores for cell lines and tumors with TP53 mutations in Machine Learning identified Cluster 1. These mutation clusters are associated with downregulated ESR1 activity. b) Boxplots showing normalized GSEA (pathway expression) and normalized CRISPR dependency scores for cell lines and tumors with TP53 mutations in Machine Learning identified Cluster 2. These mutation clusters are associated with upregulated ESR1 activity.

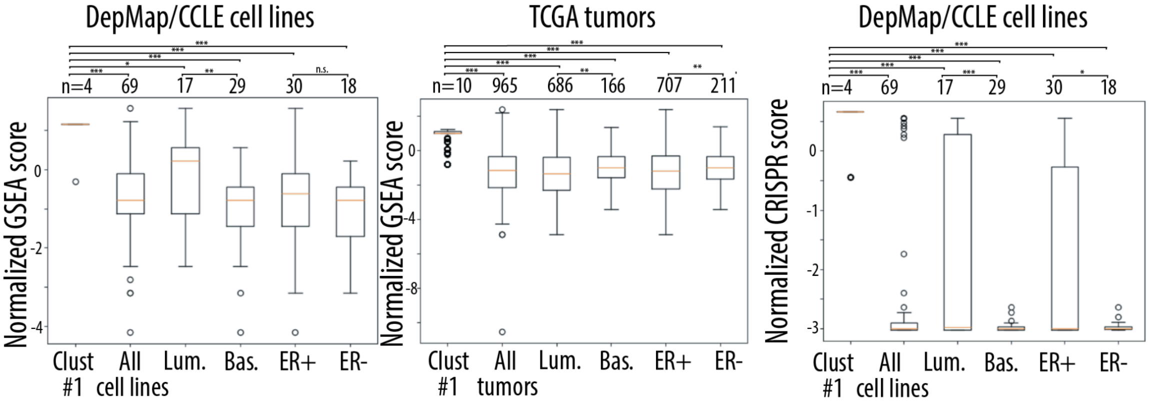


**SI 24:** a) Boxplots showing normalized GSEA (pathway expression) and normalized CRISPR dependency scores for cell lines and tumors with PTEN mutations in the Machine Learning identified Cluster. These mutation clusters are associated with downregulated ESR1 activity.

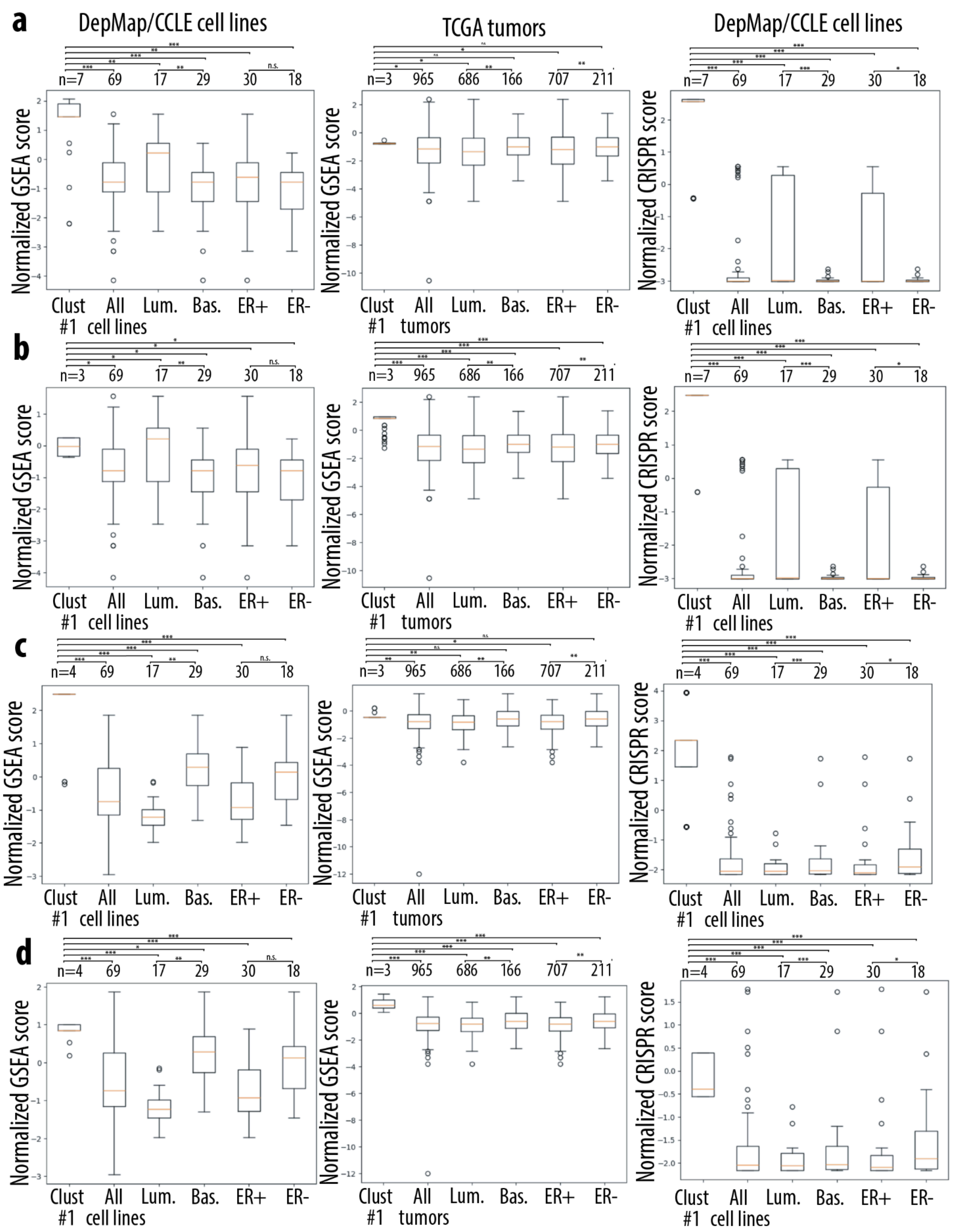


**SI 25:** a) Boxplots showing normalized GSEA (pathway expression) and normalized CRISPR dependency scores for cell lines and tumors with DST mutations in the Machine Learning identified Cluster. These mutation clusters are associated with upregulated ESR1 activity. b) Boxplots showing normalized GSEA (pathway expression) and normalized CRISPR dependency scores for cell lines and tumors with MGAM

mutations in the Machine Learning identified Cluster. These mutation clusters are associated with downregulated ESR1 activity. c) Boxplots showing normalized GSEA (pathway expression) and normalized CRISPR dependency scores for cell lines and tumors with TUBA1A mutations in the Machine Learning identified Cluster. These mutation clusters are associated with upregulated EZH2 activity. d) Boxplots showing normalized GSEA (pathway expression) and normalized CRISPR dependency scores for cell lines and tumors with KMT2C mutations in the Machine Learning identified Cluster. These mutation clusters are associated with upregulated EZH2 activity.

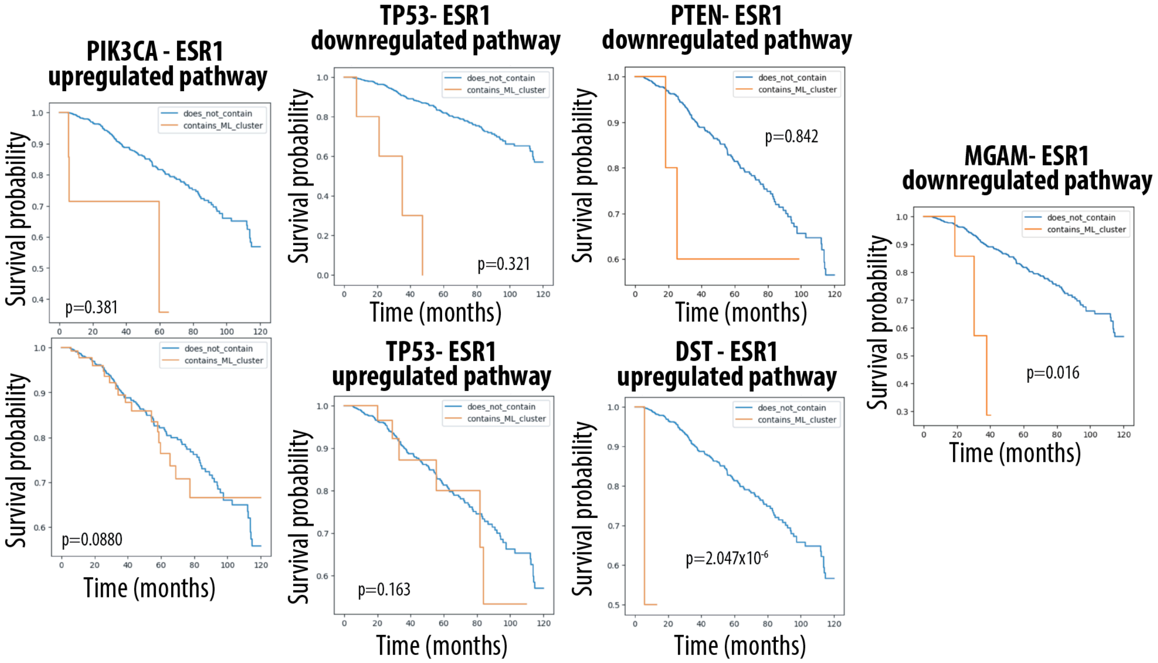


**SI 26:** TCGA survival curves for each tumor containing variants in a Machine Learning identified cluster compared to all other breast cancer tumors, shows here for associations with ESR1 pathway activity

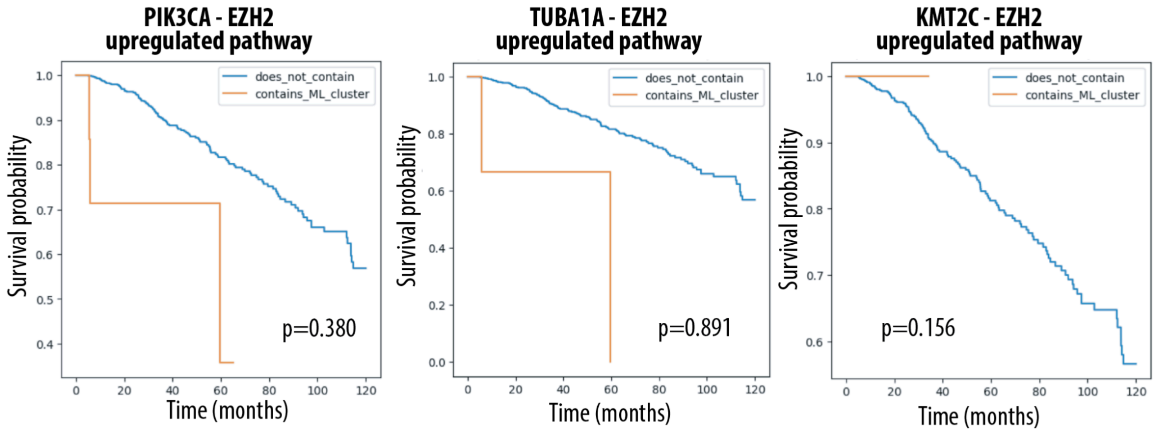


**SI 27:** TCGA survival curves for each tumor containing variants in a Machine Learning identified cluster compared to all other breast cancer tumors, shows here for associations with EZH2 pathway activity

**SI 28:** Functional impacts of each mutation cluster

| Cluster | Effect | Source |
|---|---|---|
| PIK3CA_1 | Gain of Function | BRENDA Enzyme Database, Uniprot |
| PIK3CA_2 | Gain of Function | BRENDA Enzyme Database, Uniprot |
| PIK3CA_3 | Associated with p85 dependency | BRENDA Enzyme Database, Uniprot |
| TP53_1 | Loss of Function | MAVE Database |
| TP53_2 | Loss of Function | MAVE Database |
| PTEN | Loss of Function | MAVE Database, Uniprot |
| DST | Unknown | N/A |
| MGAM | Unknown | N/A |
| KMT2C | Unknown | N/A |
| TUBA1A | Unknown | N/A |

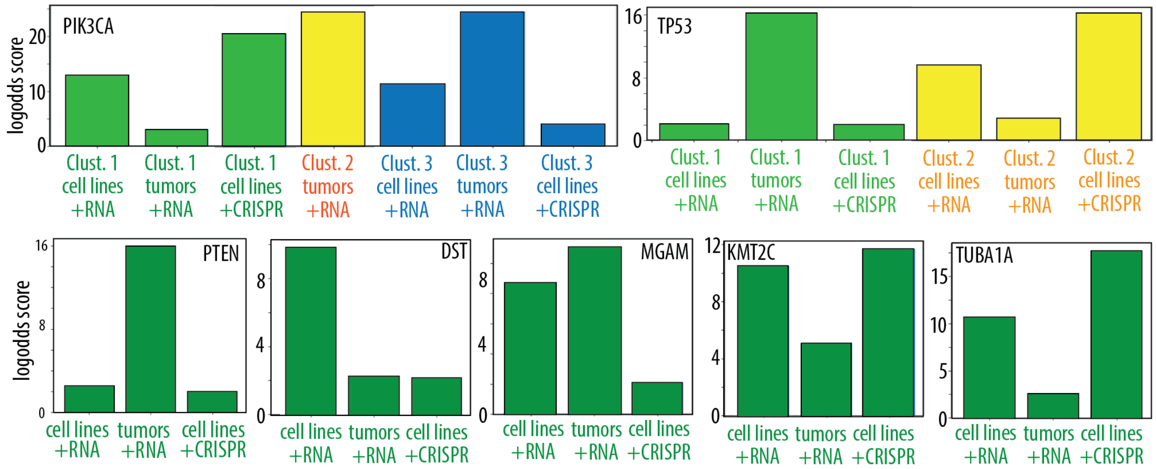


**SI 29:** Logodds plots for all clusters with all combinations of data (cell lines, tumors, RNAseq, CRISPR)

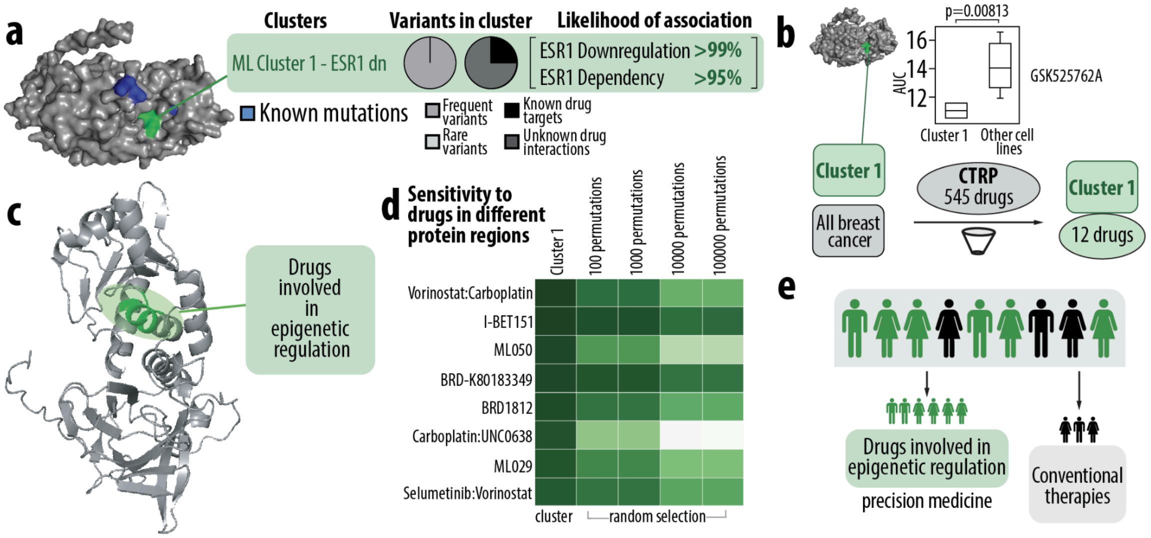


**SI 5:** a) One dense variant clusters in the PTEN protein is identified via our Machine Learning method. Cluster 1 (green) variants are associated with downregulation of the ESR1 pathway. Variants with known clinical significance are highlighted in blue. The center pie charts show the number of frequent vs rare variants and variants that are known drug targets vs variants with unknown drug interactions from Onco KB present in PTEN, and in the Machine Learning identified cluster. The likelihood that a cluster is associated with a given pathway is shown on the left. b) A schema showing the process for finding drugs with statistically different effects on cell lines containing the ML identified clusters, shown here for Cluster 1. Cell lines are sorted into two groups: those containing Cluster 1 variants, and those not containing Cluster 1 variants. Statistical analysis is performed to find all drugs that have lower AUC values for Cluster 1 cell lines and statistically different distributions compared to cell lines with no Cluster 1 mutations. c) Different regions of the protein are associated with increased sensitivity towards different drugs. Cell lines with Cluster 1 variants have greater sensitivity towards drugs involved in epigenetic regulation. d) Heat maps showing the p-values generated by the statistical analysis described in part b for Cluster 1. Additional p values from random permutation analysis are also shown. e) These results can be used to inform precision medicine treatments for patients with mutations in the indicated clusters.

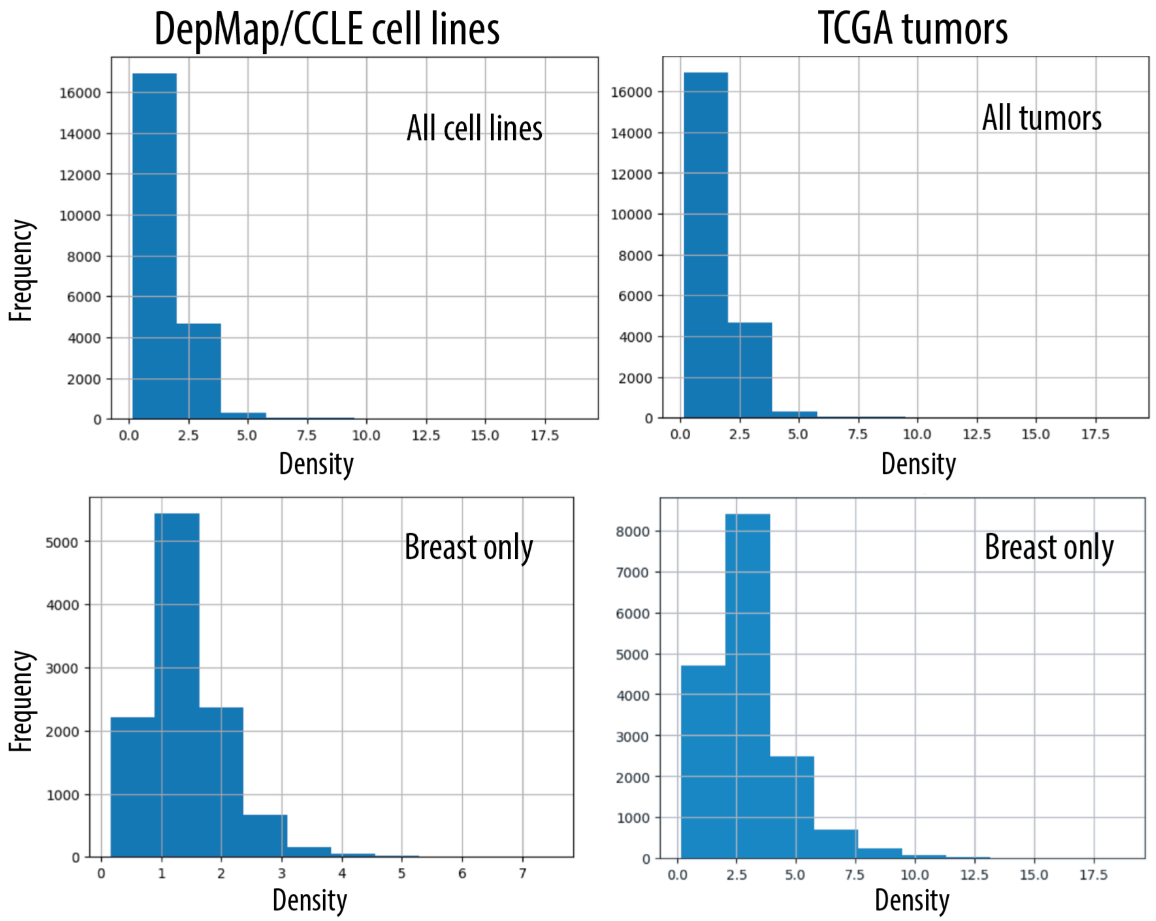


**SI 6:** Density of all clusters vs frequency of occurrence across different cell lines/tumors

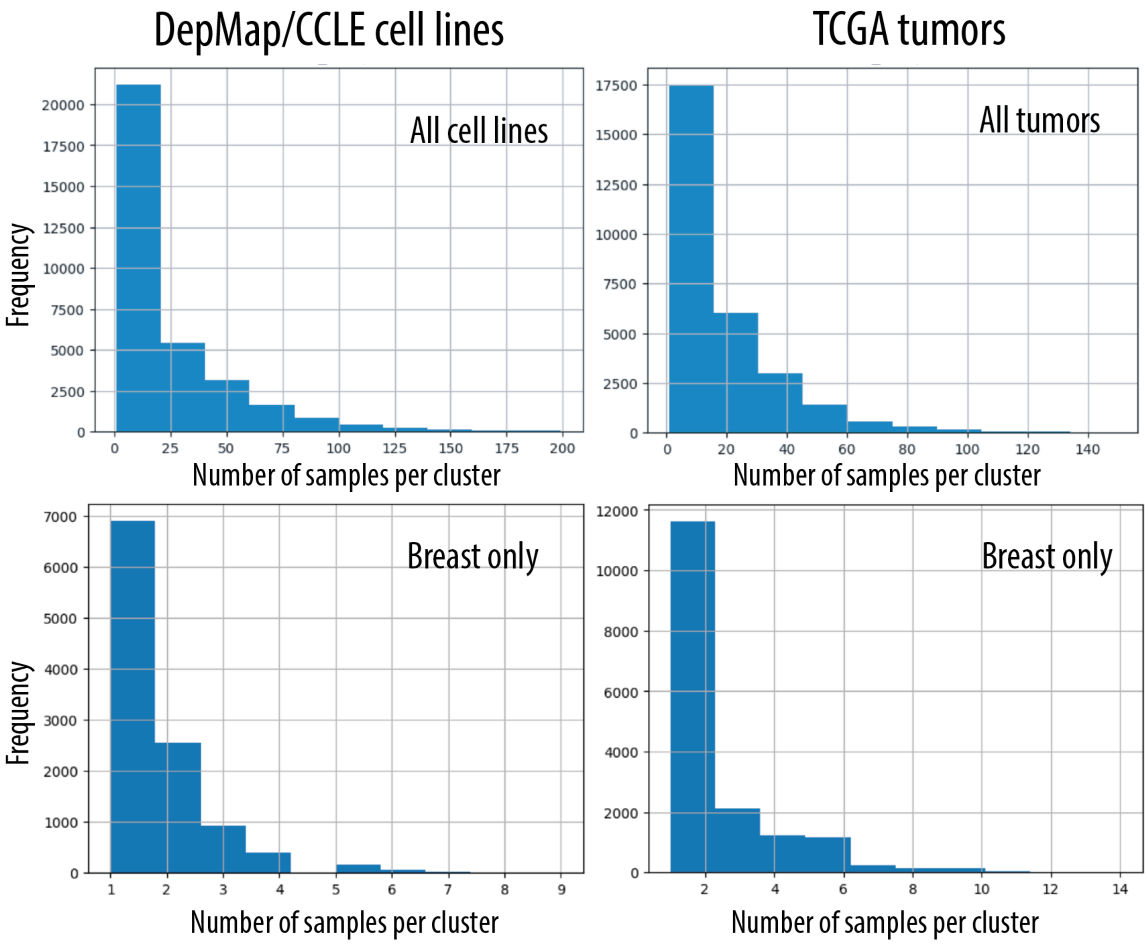


**SI 7:** Number of samples per cluster for all clusters vs frequency of occurrence across different cell lines/tumors

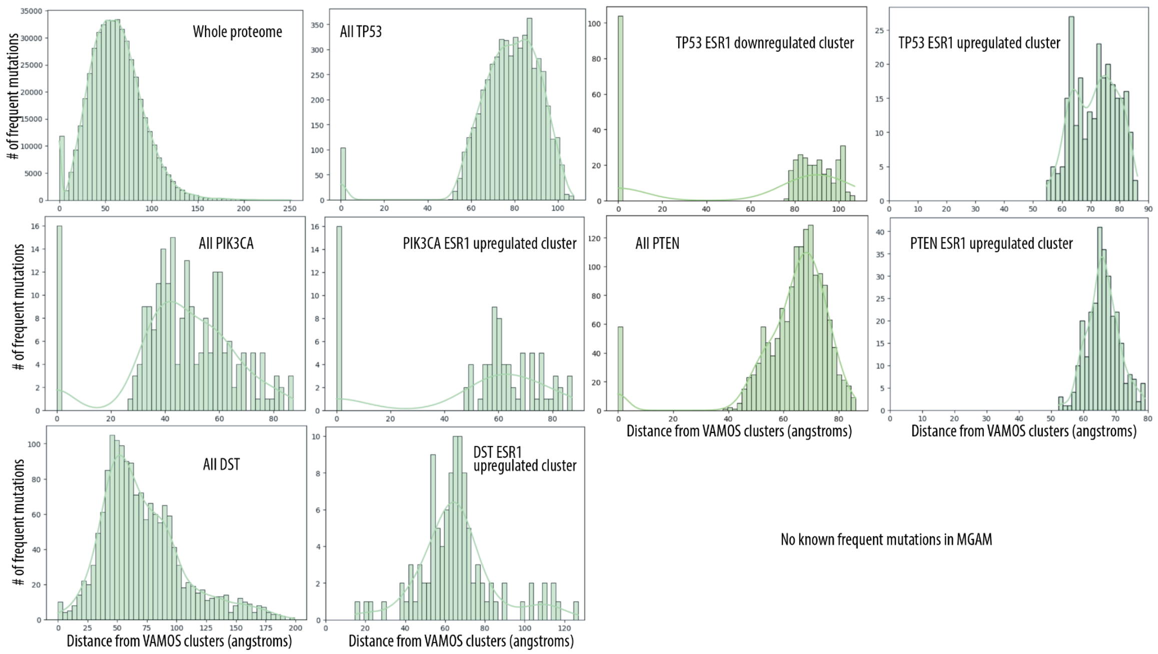


**SI 30:** Distance from known frequent mutations for each Machine learning identified cluster, shown here for clusters associated with ESR1 pathway activity

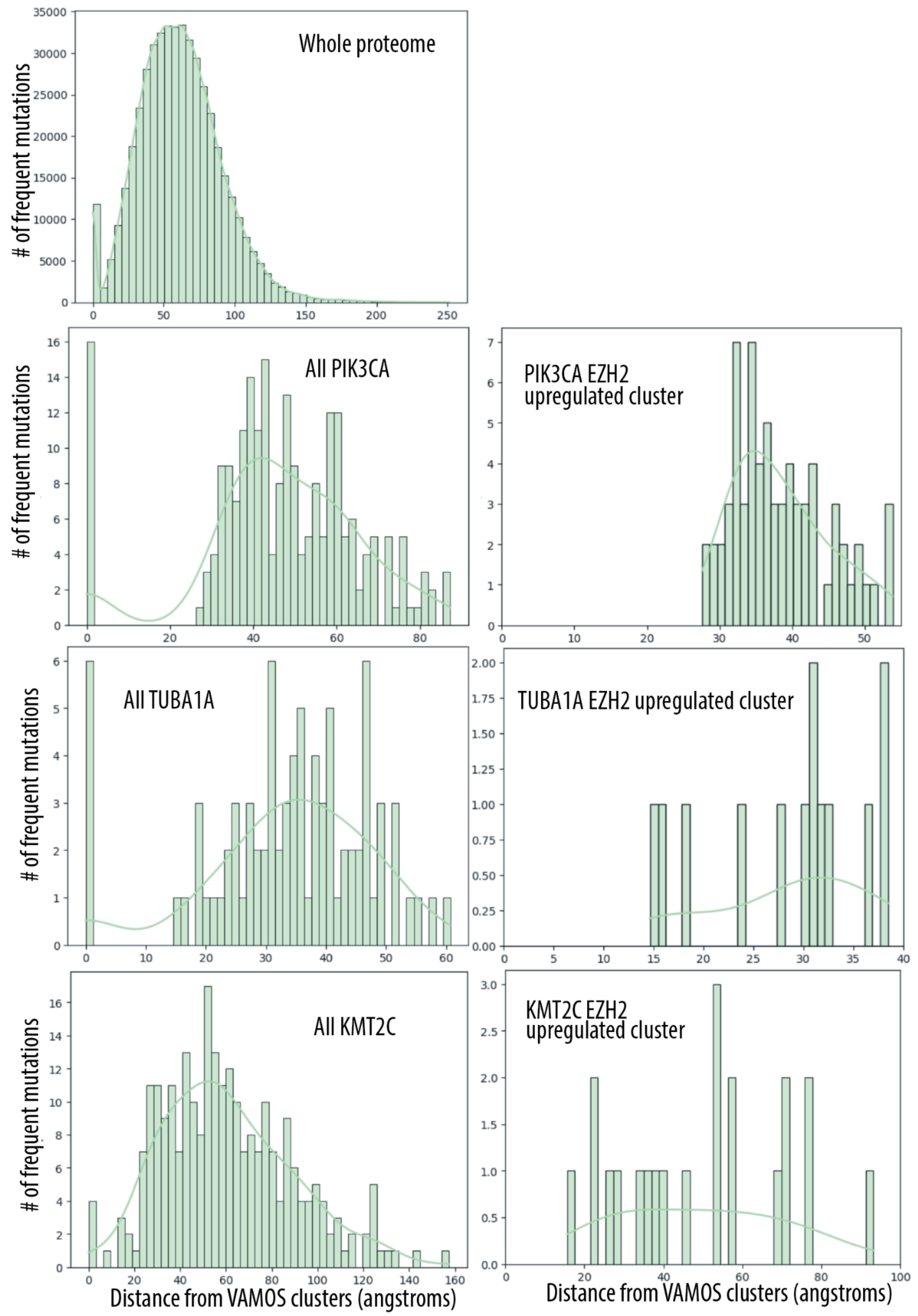


**SI 31:** Distance from known frequent mutations for each Machine learning identified cluster, shown here for clusters associated with EZH2 pathway activity

**SI 32:** Full table of proteins with their priority scores for the ESR1 pathway

| gene | priority |
|---|---|
| PIK3CA | 24.427 |
| TP53 | 23.623 |
| PTEN | 18.581 |
| KRAS | 17.387 |
| FOXA1 | 16.794 |
| HS6ST1 | 16.457 |
| ERBB2 | 15.849 |
| CHD4 | 13.998 |
| FRG1 | 13.525 |
| MUC4 | 13.284 |
| GREB1L | 13.214 |
| SF3B3 | 13.102 |
| TMEM132D | 12.934 |
| MYO1F | 12.899 |
| CPA1 | 12.821 |
| FRMPD2 | 12.543 |
| KIF1A | 11.918 |
| CD9 | 11.704 |
| KIF16B | 11.408 |
| ZNF208 | 11.184 |
| COL22A1 | 11.002 |
| KMT2C | 10.859 |
| TUBA1A | 10.840 |
| LRP1B | 10.735 |
| NTF4 | 10.645 |
| KCNH1 | 10.404 |
| ARMC3 | 10.309 |
| KMT2B | 10.309 |

| | |
|---|---|
| SRCIN1 | 10.086 |
| MT-ND4 | 9.952 |
| TUBGCP6 | 9.903 |
| ZNF423 | 9.903 |
| ITIH5 | 9.903 |
| ZNF700 | 9.798 |
| DST | 9.798 |
| CSMD2 | 9.739 |
| EP400 | 9.680 |
| LRP5 | 9.680 |
| KRT83 | 9.680 |
| MT-ND5 | 9.616 |
| GOLGB1 | 9.616 |
| MT-CYB | 9.616 |
| TCHH | 9.616 |
| MGAM | 9.190 |
| NEB | 9.105 |
| ASB10 | 8.517 |
| DNAH5 | 8.517 |
| IQGAP2 | 8.517 |
| ATP6V0A2 | 8.517 |
| BIRC3 | 8.006 |
| GGA1 | 8.006 |
| SPHKAP | 8.006 |

**SI 33:** Full table of proteins with their priority scores for the EZH2 pathway

| gene | priority |
|---|---|
| PIK3CA | 21.966 |
| PTEN | 17.285 |
| ERBB2 | 16.327 |
| FOLH1 | 15.829 |
| KRAS | 15.587 |
| FOXA1 | 15.411 |
| GPR32 | 12.660 |
| COL22A1 | 12.073 |
| MT-ND5 | 11.791 |
| KMT2C | 11.670 |
| TP53 | 11.184 |
| TUBA1A | 10.968 |
| EP400 | 10.779 |
| TRIO | 10.463 |
| LRP1B | 10.423 |
| NTNG2 | 10.309 |
| ZFHX4 | 10.138 |
| LRP5 | 10.043 |
| MUC4 | 9.999 |
| CSMD2 | 9.962 |
| KRT83 | 9.792 |
| FMN2 | 9.707 |
| EXOC4 | 9.680 |
| DST | 9.581 |
| DSE | 9.473 |
| MGAM | 9.393 |
| ZNF700 | 9.349 |
| GOLGB1 | 9.210 |

| | |
|---|---|
| FAT4 | 9.123 |
| ZNF423 | 8.987 |
| ZFYVE9 | 8.987 |
| ADCY8 | 8.987 |
| CNTNAP2 | 8.854 |
| MT-CYB | 8.854 |
| BPTF | 8.700 |
| NTF4 | 8.700 |
| RPS17 | 8.294 |
| SYNE1 | 8.294 |
| BIRC3 | 8.006 |
| IQGAP2 | 8.006 |
| HSPA12A | 8.006 |
